# Expecting the unexpected: the paranoid style of belief updating across species

**DOI:** 10.1101/2020.02.24.963298

**Authors:** Erin J. Reed, Stefan Uddenberg, Christoph D. Mathys, Jane R. Taylor, Stephanie M. Groman, Philip R. Corlett

## Abstract

Paranoia is the belief that harm is intended by others. It may arise from selective pressures to infer and avoid social threats, particularly in ambiguous or changing circumstances. We propose that uncertainty may be sufficient to elicit learning differences in paranoid individuals, without social threat. We used reversal learning behaviour and computational modelling to estimate belief updating across individuals with and without mental illness, online participants, and rats exposed to chronic methamphetamine, an elicitor of paranoia in humans. Paranoia is associated with a strong but immutable prior on volatility, accompanied by elevated sensitivity to perceived changes in the task environment. Methamphetamine exposure in rats recapitulates this impaired uncertainty-driven belief updating and rigid anticipation of a volatile environment. Our work provides evidence of fundamental, domain-general learning differences in paranoid individuals. This paradigm enables further assessment of the interplay between uncertainty and belief-updating across individuals and species.

---

Paranoia is excessive concern that harm will occur due to deliberate actions of others^1^. It manifests along a continuum of increasing severity^2-5^. Fleeting paranoid thoughts prevail in the general population^6^. A survey of over 7,000 individuals found that nearly 20% believed people were against them at times in the past year; approximately 8% felt people had intentionally acted to harm them^4^. At a national level, paranoia may fuel divisive ideological intolerance. Historian Richard Hofstadter famously described catastrophizing, context insensitive political discourse as the ‘paranoid style’:

> ***“The paranoid spokesman sees the fate of conspiracy in apocalyptic terms—he traffics in the birth and death of whole worlds, whole political orders, whole systems of human values. He is always manning the barricades of civilization. He constantly lives at a turning point.”***^***7***^

At its most severe, paranoia manifests as rigid beliefs known as delusions of persecution. These delusions occur frequently in psychotic illness, including nearly 90% of first episode patients^8^. However, paranoid beliefs are common across psychiatric and neurologic disorders, such as anxiety^9^, depression^10^, epilepsy^11^, and Alzheimer’s disease^12^. Psychostimulants elicit severe paranoid states. Methamphetamine evoked new paranoid ideation in nearly half of 274 respondents, particularly after repeated exposure (86%) or escalating dose (68%)^13^. Of those who became paranoid, the majority engaged in evasive defence strategies (hiding or fleeing), but 37% obtained weapons, and 15% attacked others. There is a clear need to better manage paranoia, and to understand and address its broader societal impact.

Paranoia has thus far defied explanation in mechanistic terms, either at the levels of behaviour or brain function. Obvious links with fear processing and social cognition, including sophisticated Game Theory driven approaches (such as the Dictator Game^14,15^) have largely re-described the phenomenon — people who are paranoid self-report difficulties with trust. Those difficulties are recapitulated in laboratory tasks that require trust^16^. However, large-scale online work with inter-personal, Game Theory motivated tasks has shown that paranoia is not driven by personal threat per se, but by negative social representations of others^14,15^. We and others have argued that such reputations are learned^17,18^, via the same fundamental learning mechanisms^19^ that stimulate non-social learning in non-human species^20^. We hypothesize that domain-general learning differences, particularly in the processing of uncertainty, underlie paranoia.

In prior work, we have shown that prediction errors, mismatches between expectation and experience that drive learning in non-human species^21^, contribute to the formation of causal beliefs and delusions in humans^22,23^. However, delusion maintenance, which we conceive of as impaired belief updating, has yet to be related definitively to specific learning mechanisms. Higher order beliefs or expectations about the noisiness of the environment may constrain whether we update beliefs or dismiss surprises as probabilistic anomalies.

Expected uncertainty, also described as risk, provides one such constraint: the perceived probabilistic variability in an environment^24^. The higher the expected uncertainty, the less surprising an atypical outcome may be, and the less compelling it is for driving belief updates. Unexpected uncertainty, in contrast, describes perceived change in the underlying statistics of the environment^25-27^. This perception promotes new learning and revision of past beliefs^24,28^. Hofstadter’s description of ‘paranoid style’ evokes the concept of unexpected uncertainty — i.e., living ‘constantly…at a turning point.’^7^ Excessive unexpected uncertainty is consistent with evolutionary theories attributing paranoia to the need to flexibly categorize or re-categorize social threats^16^. On the other hand, persecutory delusions are resistant to belief updating by definition, and even subclinical paranoia has been associated with reduced sensitivity to meaningful information in a task environment^29^.

To address this seeming paradox – excessive and deficient belief updating in paranoia – we behaviourally and computationally dissected learning mechanisms in settings of expected and unexpected uncertainty. Given our premise that paranoid learning arises from domain-general mechanisms, we invited participants to complete a non-social, three-option probabilistic learning task. Participants learn and update reward associations in response to perceived probabilistic variability of outcomes, anticipated but temporally uncertain exchange of reward probabilities between options (reversal events), and unanticipated changes in the underlying probabilities themselves (context change). This task challenges participants to update beliefs about the value of each option and the volatility of the task environment. The Hierarchical Gaussian Filter (HGF)^30,31^, a generative model of Bayesian belief, allows us to infer parameters governing learning rates from expected and unexpected variation in the task environment, initial beliefs (i.e., priors) for task volatility, and readiness to learn about changes in the task volatility itself. Beliefs concerning the values of each option update according to prediction errors weighted by belief precision; volatility prediction errors drive updates at higher levels of belief (i.e., beliefs about context). We examined the behavioural and computational correlates of paranoia both in-person and in a large online sample, spanning patients and healthy controls with varying degrees of paranoia. We also undertook a pre-clinical replication in rodents exposed chronically to saline or methamphetamine^32^. We predicted that paranoia-related learning differences would be particularly prominent in settings of contextual change. We observed elevated sensitivity to unexpected uncertainty resulting in excessive revision of option-outcome associations, accompanied by elevated volatility priors and deficient learning about contextual change (metavolatility).

## Results

We analysed belief updating across three reversal-learning experiments (Fig. 1): an in laboratory pilot of patients and healthy controls, stratified by stable, paranoid personality trait (Experiment 1); four online task variants administered to participants via the Amazon Mechanical Turk (MTurk) marketplace (Experiment 2); and a re-analysis of data from rats on chronic, escalating doses of methamphetamine, a translational model of paranoia (Experiment 3)^32^.

**Fig. 1.**
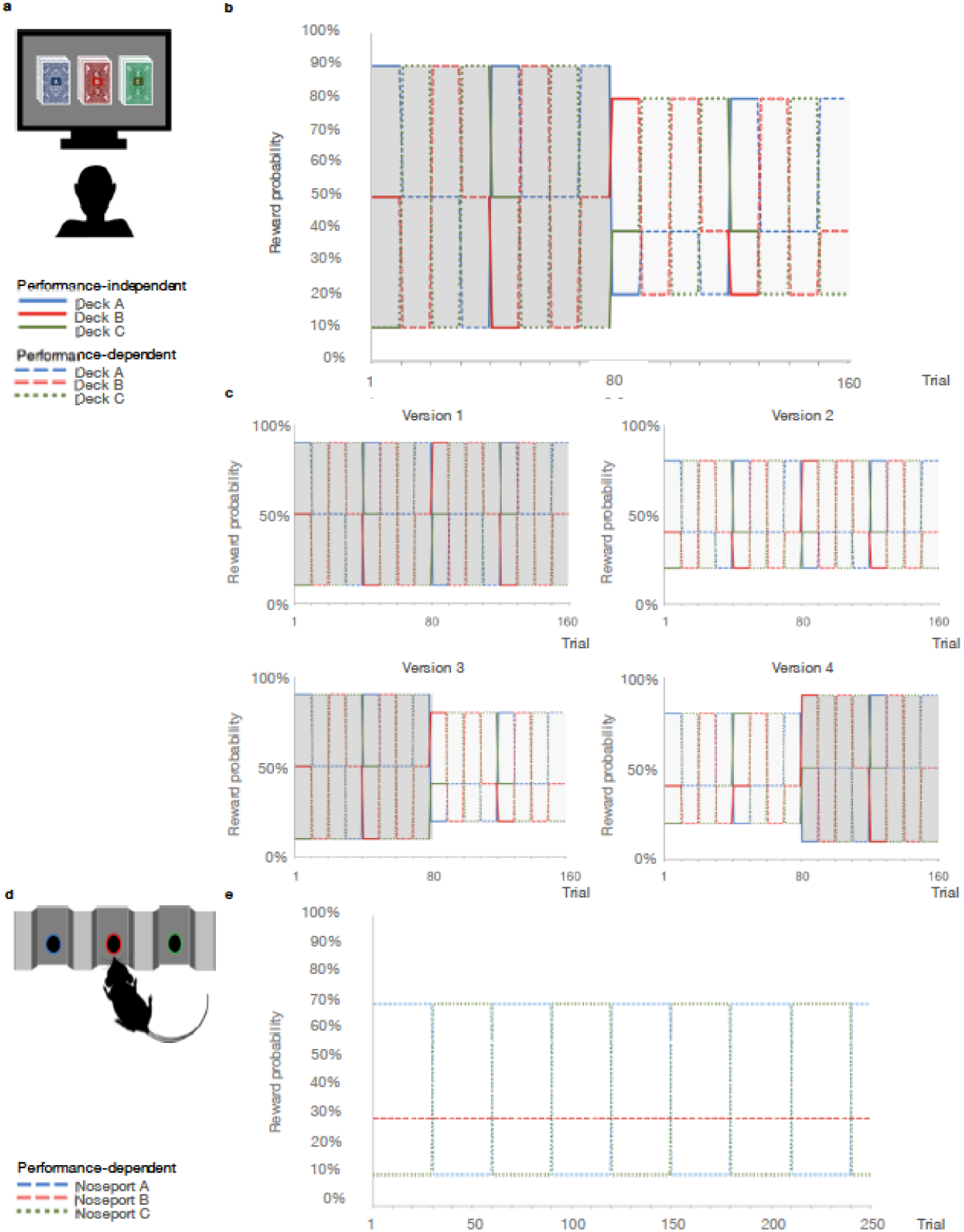
Probabilistic reversal learning task. **a**, Human paradigm: participants choose between three ecks of cards with different, unknown probabilities of reward and loss. **b**, Reward contingency schedule for in laboratory experiment. On trial 81, the probability context shifts from 90%, 50%, and 10% (dark grey) to 80%, 40%, and 20% without warning (light grey). **c**, Reward contingency schedules for online experiment. **d**, Rat paradigm: subjects choose between three noseports with different probabilities of sucrose pellet reward. **e**, Reward contingency schedule for rat experiment^39^.

**Experiment 1.** First, we explored trans-diagnostic associations between paranoia and performance on a reversal-learning paradigm. Participants (*n*=32) with and without psychiatric diagnoses (anxiety, depression, bipolar disorder, schizophrenia, and schizoaffective disorder) completed questionnaire versions of the *Structured Clinical Interview for DSM-IV Axis II Personality Disorders* (SCID-II) screening assessment^33^, Beck’s Anxiety Inventory (BAI)^34^, Beck’s Depression Inventory (BDI)^35^, and demographic assessments (Table 1). Approximately two-thirds of participants endorsed three or fewer items on the SCID-II paranoid personality subscale (median=1 item). Participants who endorsed four or more items were classified as high paranoia (*n*=11), consistent with the diagnostic threshold for paranoid personality disorder. Low paranoia (*n*=21) and high paranoia groups did not differ significantly by age, nor were there significant group associations with gender, educational attainment, ethnicity, or race, although a larger percentage of paranoid participants identified as racial minorities or “not specified” (Table 1). Diagnostic category (i.e., healthy control, mood disorder, or schizophrenia spectrum) was significantly associated with paranoia group membership, χ^2^ (2, *n=*32)=12.329, *P=*0.002, Cramer’s V=0.621, as was psychiatric medication usage, χ^2^ (1, *n=*32)=9.871, *P*=0.003, Cramer’s V=0.555. These differences were due largely to the higher proportion of healthy controls in the low paranoia group. SCID-II paranoia scores correlated with symptoms of anxiety and depression (BAI: Pearson’s *r*=0.611, *P*=0.0002, 95% Confidence Interval (CI)=[0.315,0.906]; BDI: Pearson’s *r*=0.564, *P*=0.001, CI=[0.257, 0.872]). As expected, paranoia, BAI, and BDI scores were significantly elevated in the high paranoia group relative to low paranoia controls (Table 1; paranoia: mean difference (MD)=0.536, CI=[0.455,0.618], *t*(30)=13.476, *P*=2.92E-14, Hedges’ *g*=5.016; BAI: MD=0.585, CI=[0.239, 0.931], *t*(30)=3.453, *P*=0.002, Hedges’ *g*=1.285, MD=-0.585; BDI: MD=0.427, CI=[0.078, 0.775], *t*(11.854)=2.67, *P*=0.021, Hedges’ *g*=1.255).

**Table 1.**
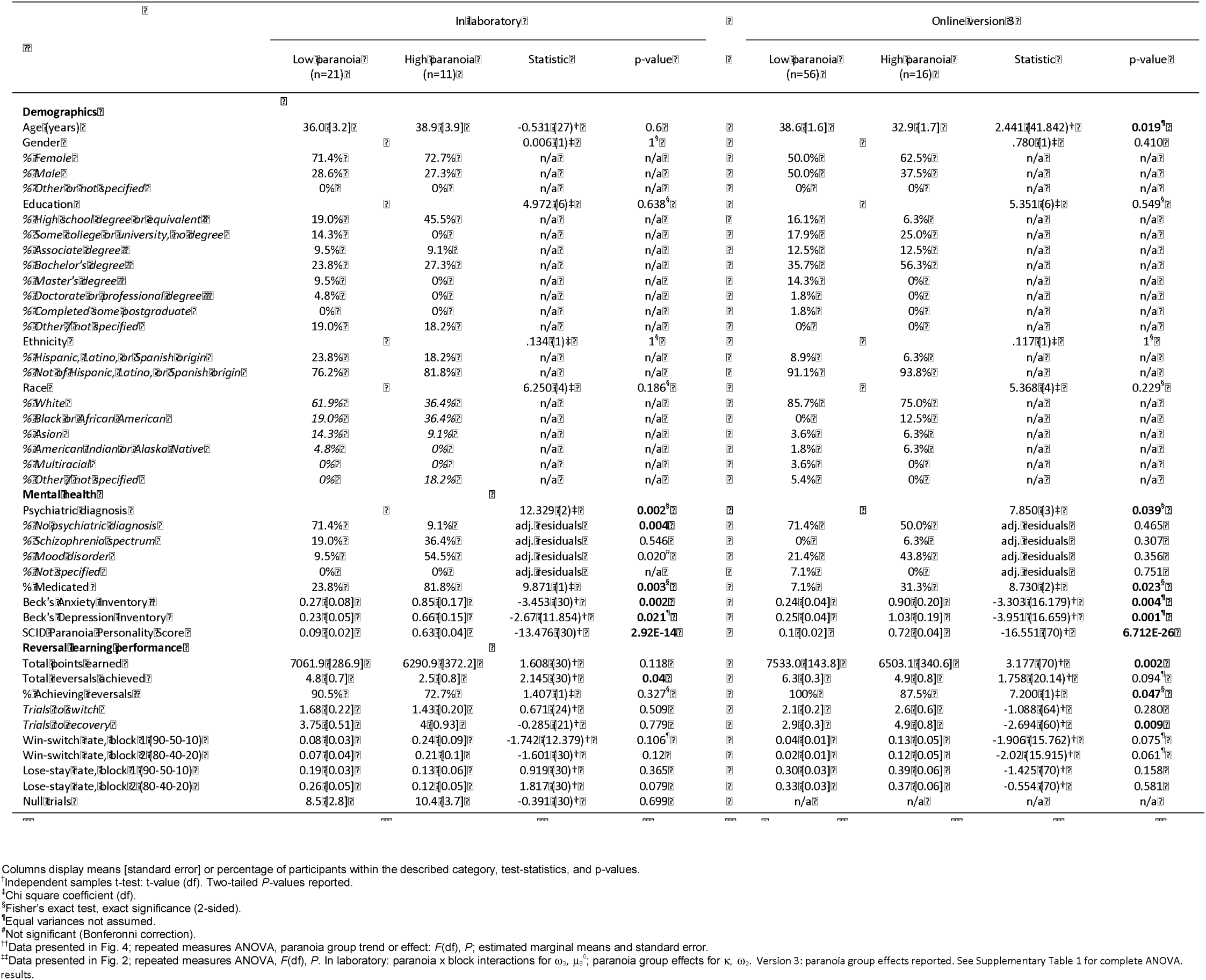
In laboratory vs. online version 3

Participants completed a three-option reversal-learning task in which they chose between three decks of cards with hidden reward probabilities (Fig. 1 a and b). They selected a deck on each turn and received positive or negative feedback (+100 or −50 points, respectively). They were instructed to find the best deck with the caveat that this deck may change. Undisclosed to participants, reward probabilities switched among decks after selection of the highest probability option in nine out of ten consecutive trials (“reversal events”). Reward probability context changed from 90%, 50%, and 10% chance of reward to 80%, 40%, and 20% between the first and second halves of the task (“contingency context change”; block 1=80 trials, 90-50-10%; block 2=80 trials, 80-40-20%). High paranoia subjects achieved fewer reversals (MD=-2.31, CI=[-4.504, −0.111,], *t*(30)=-2.145, *P*=0.04, Hedges’ *g*=0.798), but total points earned did not significantly differ (Table 1).

**Experiment 2.** We replicated the effects of paranoia on reversal-learning in a larger online sample. We also tested alternative task versions to control for the contingency context change (Fig. 1c). Version 1 (*n*=45 low paranoia, 20 high paranoia) provided a constant contingency context of 90-50-10% reward probabilities; version 2 (*n*=69 low paranoia, 18 high paranoia) provided a constant context of 80-40-20%; version 3 (*n*=56 low paranoia, 16 high paranoia) replicated Experiment 1 with a context change from 90-50-10% to 80-40-20%; version 4 (*n*=64 low paranoia, 19 high paranoia) provided the reverse context change, 80-40-20% to 90-50-10%. Demographic and mental health questionnaire responses did not differ significantly across task versions (Table 2). Total points and reversals achieved suggest variations in task difficulty (Table 2, version effects: points earned, *F*(3)=232.88, *P*=4.16E-18, η_p_^2^=0.245; reversals achieved, *F*(3)=4.329, *P*=0.005, η_p_^2^=0.042), but there was no significant association between task version and attrition rate (52.7%, 52.9%, 54.6%, and 53.1% attrition, respectively; χ^2^(3)=0.167, *P*=0.983, Cramer’s V=0.015).

**Table 2.**
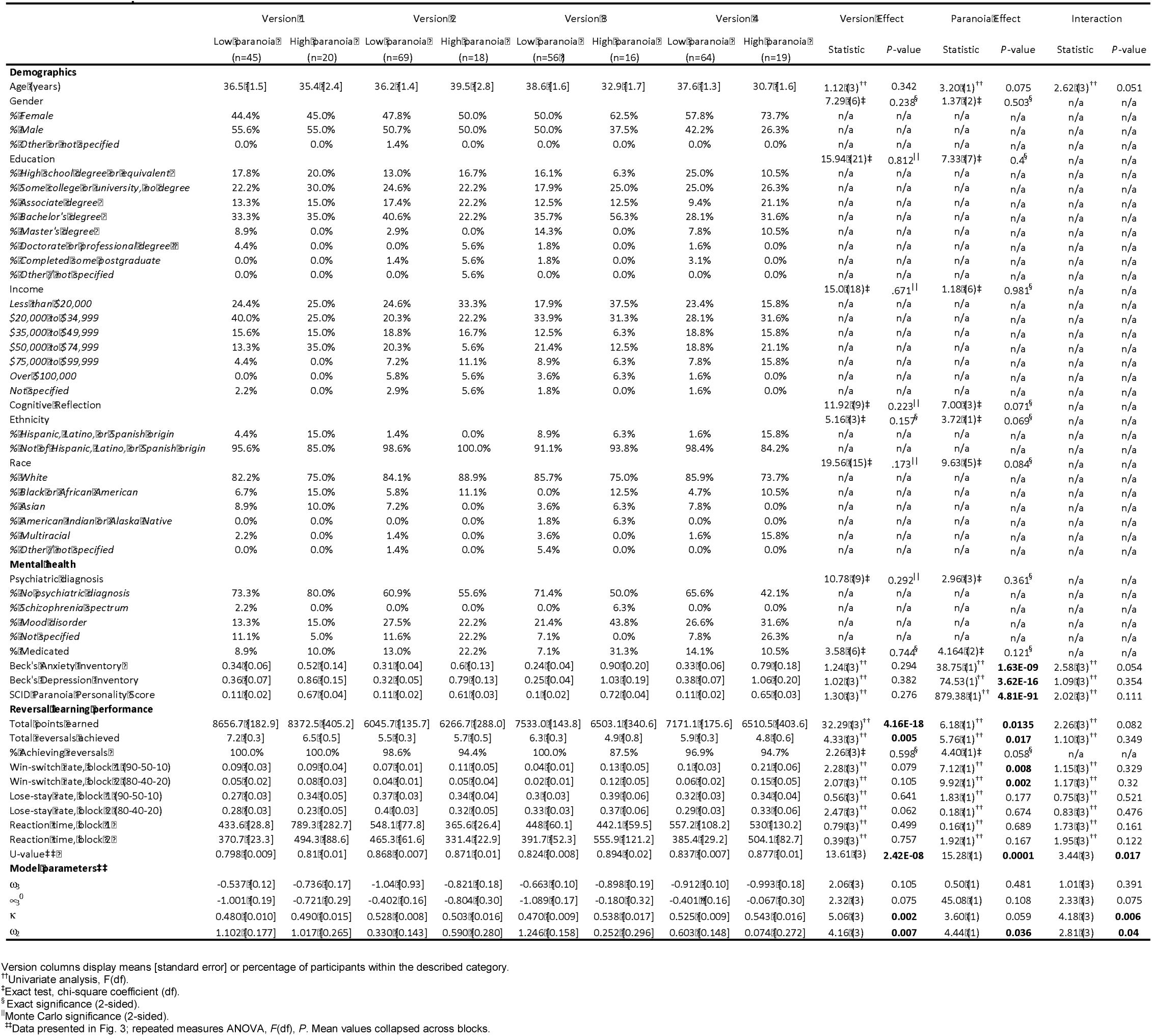
Online experiment

Across task versions, high paranoia participants endorsed higher BAI and BDI scores (*n*=73 high paranoia, 234 low paranoia; BAI: *F*(1)=38.752, *P*=1.63E-09, η_p_^2^=0.115; BDI: *F*(1)=74.528, *P*=3.62E-16, η_p_^2^=0.20; Table 2). Both correlated with paranoia (BAI: Pearson’s *r*=0.450, *P*=1.09E-16, CI=[0.348, 0.55]; BDI: Pearson’s *r*=0.543, *P*=6.26E-25, CI=[0.448, 0.638]). Trial-by-trial reaction time did not differ significantly between low and high paranoia (Table 2), but high paranoia participants earned fewer total points (*F*(1)=6.175, *P*=0.014, η_p_^2^=0.020) and achieved fewer reversals (*F*(1)=5.762, p=0.017, η_p_^2^=0.019; Table 2). Deck choice perseveration after negative feedback (lose-stay behaviour) did not significantly differ by paranoia group, but choice switching after positive feedback (win-switch behaviour) was elevated in high paranoia (block 1: *F*(1)=7.117, *P*=0.008, η_p_^2^=0.023; block 2: *F*(1)=9.918, *P*=0.002, η_p_^2^=0.032; Table 2).

**Experiment 3.** To translate across species, we performed a new analysis of published data from rats exposed to chronic methamphetamine^32^. Rats chose between three operant chamber noseports with differing probabilities of sucrose reward (70%, 30%, and 10%; Fig.1 d and e). Contingencies switched between the 70% and 10% noseports after selection of the highest reinforced option in 21 out of 30 consecutive trials (Fig. 1e). Rats were tested for 26 within-session reversal blocks (Pre-Rx, *n*=10 per group), administered saline or methamphetamine according to a 23-day schedule mimicking the escalating doses and frequencies of chronic human methamphetamine users^32^, and tested once per week for four weeks following completion of the drug regimen (Post-Rx; *n*=10 saline, 7 methamphetamine)^32^. Relative to rats exposed to saline, those rats exposed to methamphetamine exhibited increased win-switch behaviour and perseveration after negative feedback^32^.

### Computational modelling

We employed hierarchical Gaussian filter (HGF) modelling to compare belief updating across individuals with low and high paranoia, as well as across human participants and rats exposed to methamphetamine (Table 3). We paired a three-level perceptual model with a softmax decision model dependent upon third level volatility (Fig. 2a). We inverted the model from subject data (trial-by-trial choices and feedback) to estimate parameters for each individual (Fig. 2b). Level 1 (x_1_) characterizes trial-by-trial perception of task feedback (win or loss in humans, reward or no reward in rats), Level 2 (x_2_) distinguishes stimulus-outcome associations (deck or noseport values), and Level 3 (x_3_) renders perception of the overall reward contingency context (i.e., volatility or variance of the deck or noseport values). Belief trajectories were unique to each subject due to the probabilistic, performance-dependent nature of the task, but we estimated initial beliefs (priors) for x_2_ and x_3_ (μ_2_^0^ and μ_3_^0^, respectively). We also estimated ω_2,_ the contribution of tonic (expected) volatility on learning stimulus-outcome associations, and κ, the coupling or impact of phasic (unexpected) volatility (x_3_) on the x_2_ belief trajectory. In the setting of our tasks, these two parameters best capture the effects of expected and unexpected uncertainty in updating stimulus-outcome associations. Higher coupling (κ) implies faster belief updating in response to perceived change at the level above, whereas lower or more negative values suggest slower updating. Diminished ω_2_ indicates more rigid beliefs about the underlying risk or probability of each option. The third parameter, ω_3,_ characterizes perception of ‘metavolatility,’ the tonic volatility of the volatility itself (i.e., how stable the changes in underlying contingencies of the decks might be)^36^. The lower ω_3_ is, the slower a subject is to update beliefs about contextual volatility.

**Table 3.**
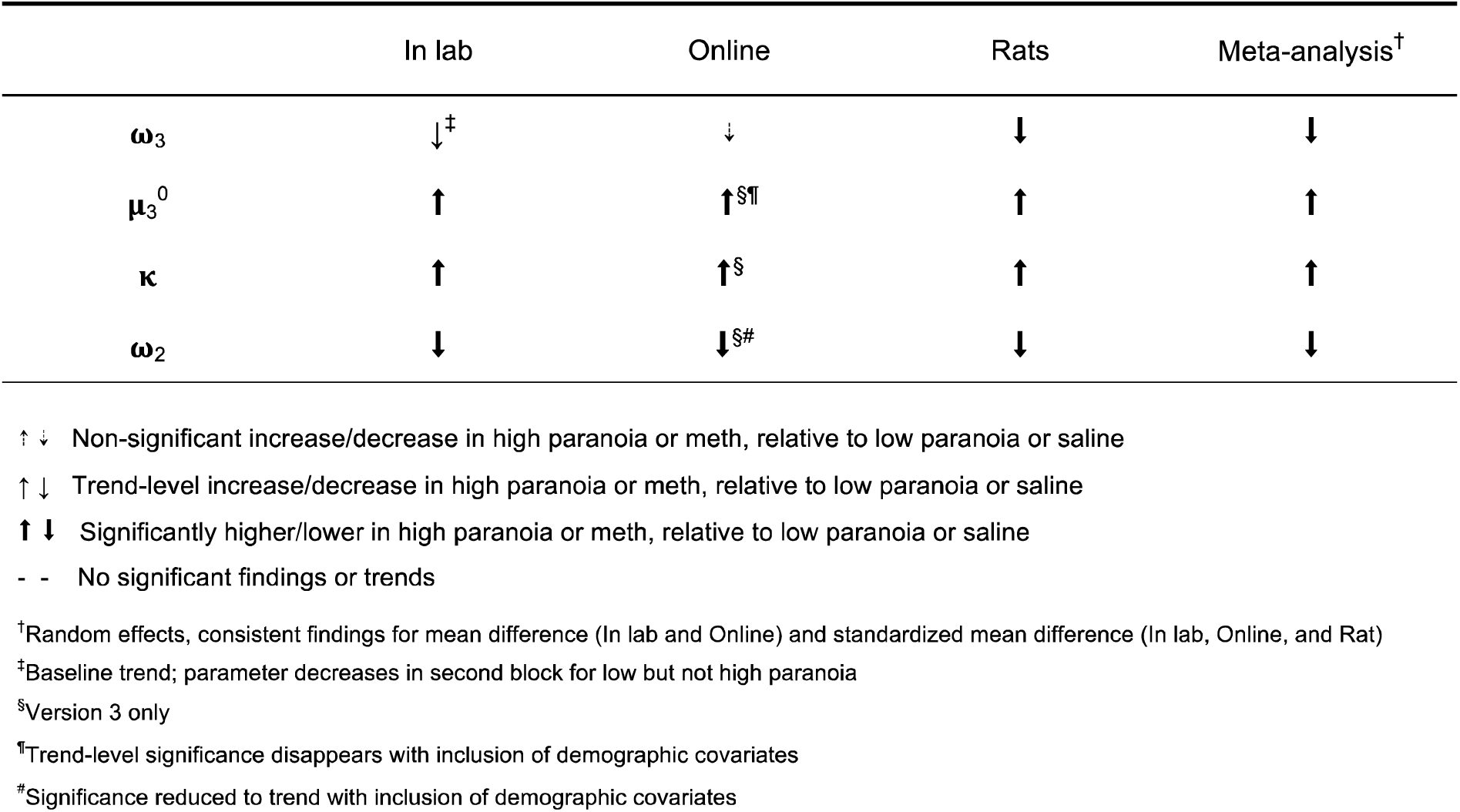
Summary of Paranoia/Methamphetamine Effects on Belief-Updating.

**Fig. 2.**
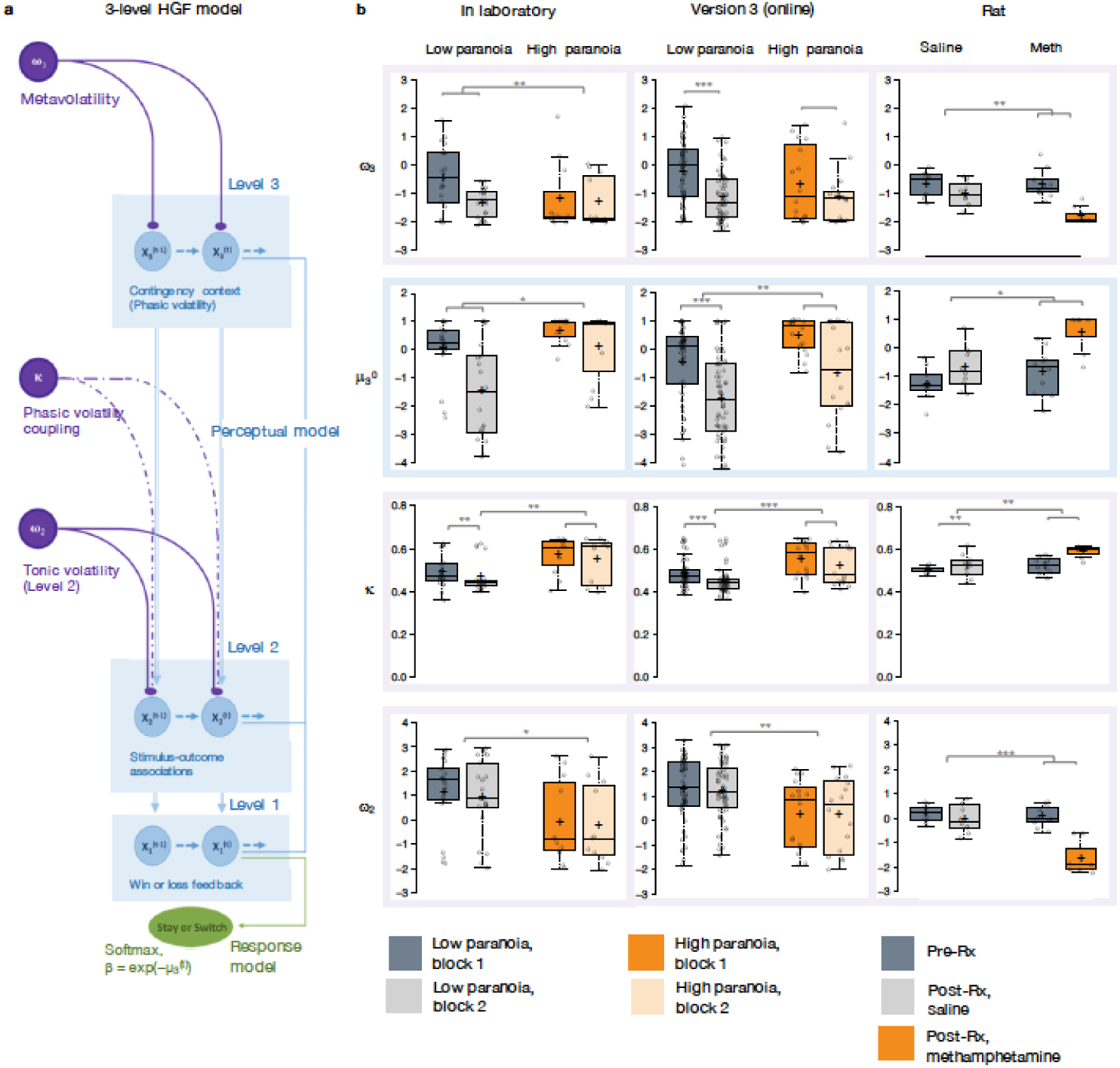
Hierarchical Gaussian Filter (HGF) model parameters. **a**, 3-level HGF perceptual model (blue) with a softmax decision model (green). Level 1 (x_1_) corresponds to trial-by-trial perception of win or loss feedback. Level 2 (x_2_) represents stimulus-outcome associations (e.g., deck values). Level 3 (x_3_) models perception of the overall reward contingency context. The impact of phasic volatility upon x_2_ is captured by κ, the coupling parameter. Tonic volatility modulates x_3_ and x_2_ via ω_3_ and ω_2_, respectively. μ_3_^0^ is the initial value of the third level volatility belief. **b**, Estimated parameters replicate across high paranoia groups (orange) in the in laboratory experiment (n=21 low paranoia, 11 high paranoia); analogous online task (version 3, n=56 low paranoia, 16 high paranoia); and rats exposed to chronic, escalating methamphetamine (n=10 per group Pre-Rx; n=10 saline, 7 methamphetamine Post-Rx). Error bars denote standard error (SEM); *p ≤ 0.05, **p ≤ 0.01, ***p ≤ 0.001.

Priors did not differ between groups at x_2_ (Supplementary Table 1) but paranoid individuals and rats exposed to methamphetamine exhibited elevated μ_3_^0^, an initial perception of higher contingency context volatility (Fig. 2b, blue). In Experiment 1, we observed an interaction between task block and paranoia group (*F*(1)=5.344, *P*=0.028, η_p_^2^=0.151; Table 1). μ_3_^0^ differed between high and low paranoia in both blocks (block 1, *F*(1)=4.232, *P*=0.048, η_p_^2^=0.124, MD=0.658, CI=[0.005,1.312]; block 2, F(1)=7.497, *P*=0.010, η_p_^2^=0.20, MD=1.598, CI=[0.406, 2.789]), but only low paranoia subjects significantly updated their priors between block 1 and block 2 (*F*(30)=39.841, *P*=5.85E-07, η_p_^2^=0.570, MD=1.504, CI=[1.017, 1.99]). In Experiment 2, the analogous task design (version 3) demonstrated significant effects of block (*F*(1)=64.652, *P*=1.54E-11, η_p_^2^=0.480, MD=1.303, CI=[0.980,1.627]) and paranoia (*F*(1)=6.366, *P*=0.014, η_p_^2^=0.083, MD=0.909, CI=[0.191, 1.628]; Table 1). Rats showed a similar effect following methamphetamine exposure with a significant time (Pre-Rx, Post-Rx) by treatment (methamphetamine, saline) interaction (*F*(1)=5.159, *P*=0.038, η_p_^2^=0.256; pre versus post methamphetamine effect: *F*(15)=12.186, *P*=0.003, MD=1.265, CI=[-0.493, 2.037]; Pre-Rx mean [standard error]*=* −1.25 [0.56] saline, −0.77 [0.80] methamphetamine; Post-Rx: *m*=-0.69 [0.74] saline, 0.58 [0.73] methamphetamine). Random effects meta-analyses confirmed significant cross-experiment replication of elevated μ_3_^0^ in human participants with paranoia (in laboratory and online version 3; MD_META_= 1.110, CI=[0.927, 1.292], *z*_META_=11.929, p=8.356E-33) and across humans with paranoia and rats exposed to methamphetamine (MD_META_=2.090, CI=[0.123, 4.056], *z*_META_=2.083, p=0.037).

Paranoid participants and methamphetamine exposed rats updated stimulus-outcome associations more strongly in response to perceived phasic volatility (e.g., correctly or incorrectly inferred reversals; Fig. 2b). κ showed significant paranoia group and block effects across the in laboratory experiment and online version 3 (Table 1; paranoia effects, in laboratory: *F*(1)=7.599, *P*=0.010, η_p_^2^=0.202, MD=0.081, CI=[0.021, 0.140]; online version 3: *F*(1)=13.521, *P*=0.0005, η_p_^2^=0.162, MD=0.068, CI=[0.031-0.104]; MD_META_ = 0.079, CI=[0.063, 0.095], *z*_META_=9.502 p=2.067E-21); see Supplementary Table 1 for block effects). κ increased from baseline in rats on methamphetamine, yielding significant effects of treatment (*F*(1)=13.356, *P*=0.002, η_p_^2^=0.471, MD=0.045, CI=[0.019, 0.072]) and time (*F*(1)=9.132, *P*=0.009, η_p_^2^=0.378, MD=0.041, CI=[0.012, 0.069]); however, the interaction between time and treatment did not reach statistical significance (Supplementary Table 1; Pre-Rx *m=*0.499 [0.015] saline, 0.523 [0.040] methamphetamine; Post-Rx: *m=*0.518 [0.053] saline, 0.585 [0.029] methamphetamine). Replication of group effects was significant across all three experiments (MD_META_=2.063, CI=[0.341, 3.785], *z*_META_=2.348, p=0.019).

Tonic volatility and metavolatility (ω_2,_ ω_3_) were decreased in paranoid participants and rats exposed to methamphetamine (Fig. 2b). In laboratory and online (version 3), paranoid individuals were slower to update stimulus-outcome associations in response to expected stochastic variance within the contingency context (Table 1; ω_2_ paranoia effect, in laboratory: *F*(1)=4.186, *P=*0.050, η_p_^2^=0.122, MD=-1.188, CI=[-2.375, −0.002]; online version 3: *F*(1)=8.7, *P*=0.004, η_p_^2^=0.111, MD=-0.993, CI=[-1.665, −0.322]; MD_META_=-1.154, CI=[-1.455, −0.853], *z*_META_=-7.521, p=5.450E-14). The effects of methamphetamine exposure in rats were consistent (MD_META_=-1.992, CI=[-3.318, −0.665], *z*_META_=-2.943, p=0.003) yet more striking, with a strongly negative ω_2_ accounting for the more pronounced lose-stay behaviour in rats (time by treatment interaction, *F*(1)=18.454, *P*=0.001, η_p_^2^=0.552; pre versus post methamphetamine: *F*(1)=42.242, *P*=1.0E-5^32^, η_p_^2^=0.738, MD=-1.604, CI=[-2.130, −1.078]; Pre-Rx *m*=0.198 [0.33] saline, −0.036 [0.42] methamphetamine; Post-Rx: *m*=-0.023 [0.56] saline, −1.640 [0.71] methamphetamine). Metavolatility (ω_3_) was similarly lower across paranoia and methamphetamine exposed groups (in laboratory, online version 3, and rats: MD_META_=-1.155, CI=[-2.139, - 0.171], *z*_META_=-2.3, p=0.021), suggesting resistance to updating beliefs about the overall contingency context. In laboratory, we observed a block by paranoia group interaction (Table 1, *F*(1)=6.948, *P*=0.010, η_p_^2^=0.188). Post-hoc tests differentiated first and second blocks for the low paranoia group only (*F*(1)=26.640, *P*=1.5E-5, η_p_^2^=0.470, MD=-0.876, CI=[-1.222, −0.529]). The paranoia effect did not reach statistical significance for online version 3 (block effect only, *F*(1)=14.932, *P*=0.0002, η_p_^2^=0.176, MD=-0.692, CI=[-1.050, −0.335]; Supplementary Table 1), but meta-analytic random effects analysis confirms a significant paranoia group difference (in laboratory and online version 3: MD_META_=-0.341, CI=[-0.522, −0.159], *z*_META_=-3.68, p=0.0002). Methamphetamine exposure decreased ω_3_ in rats (time by treatment interaction, (*F*(1)=9.058, *P*=0.009, η_p_^2^=0.376; pre versus post methamphetamine: *F*(1)=30.668, P=5.7E-5, η_p_^2^=0.672, MD=-1.210, CI=[-1.676, - 0.745]; Pre-Rx m=-0.692 [0.44] saline, −0.607 [0.51] methamphetamine; Post-Rx: *m=*-1.044 [0.44] saline, −1.817 [0.32] methamphetamine).

We applied False Discovery Rate (FDR) corrections for modelling parameters. κ group effects survived corrections within each experiment (Supplementary Table 2). In addition to κ, μ_3_^0^ survived for experiment 1; μ_3_^0^ and ω_2_ survived in online version 3; and μ_3_^0^, ω_2_, and ω_3_ survived in experiment 3 as group effects. Such correction is not yet standard practice with this modelling approach^36-38^ but we believe it should be, and when effects survive correction we should increase our confidence in them.

### Paranoia effects across task versions

To examine the relationship between contingency context change and paranoia within our HGF parameters, we performed split-plot, repeated measures ANOVAs across all four task versions (Experiment 2; paranoia group and task version between subject factors). Paranoia group effects were specific to versions of the task in which we explicitly manipulated uncertainty via context change (Fig. 3, Supplementary Table 2). Specifically, we observed paranoia by version interactions for κ (*F*(3)=4.178, *P*=0.006, η_p_^2^=0.040) and ω_2_ (*F*(3)=2.809, *P*=0.040, η_p_^2^=0.027; Table 2). Post-hoc tests confirmed that significant paranoia group effects were restricted to version 3 (κ: *F*(1)=12.230, *P*=0.001, η_p_^2^=0.039, MD=0.068, CI=[0.03,0.106]; ω_2_: *F*(1)=8.734, *P*=0.003, η_p_^2^=0.028, MD=-0.993, CI=[-1.655, −0.332]) and a trend for version 4 (ω_2_: *F*(1)=2.909, *P*=0.089, η_p_^2^=0.010, MD=-0.528, CI=[-1.138, 0.081], Fig. 3a). μ_3_^0^ also exhibited a paranoia by version trend (Table 2, *F*(3)=2.329, *P*=0.075, η_p_^2^=0.023), largely driven by version 3 (*F*(1)=6.206, *P*=0.013, η_p_^2^=0.020, MD=0.909, CI=[0.191, 1.628]; Fig. 3a). There were no significant paranoia effects or interactions for ω_3_ (Supplementary Table 2).

**Fig. 3.**
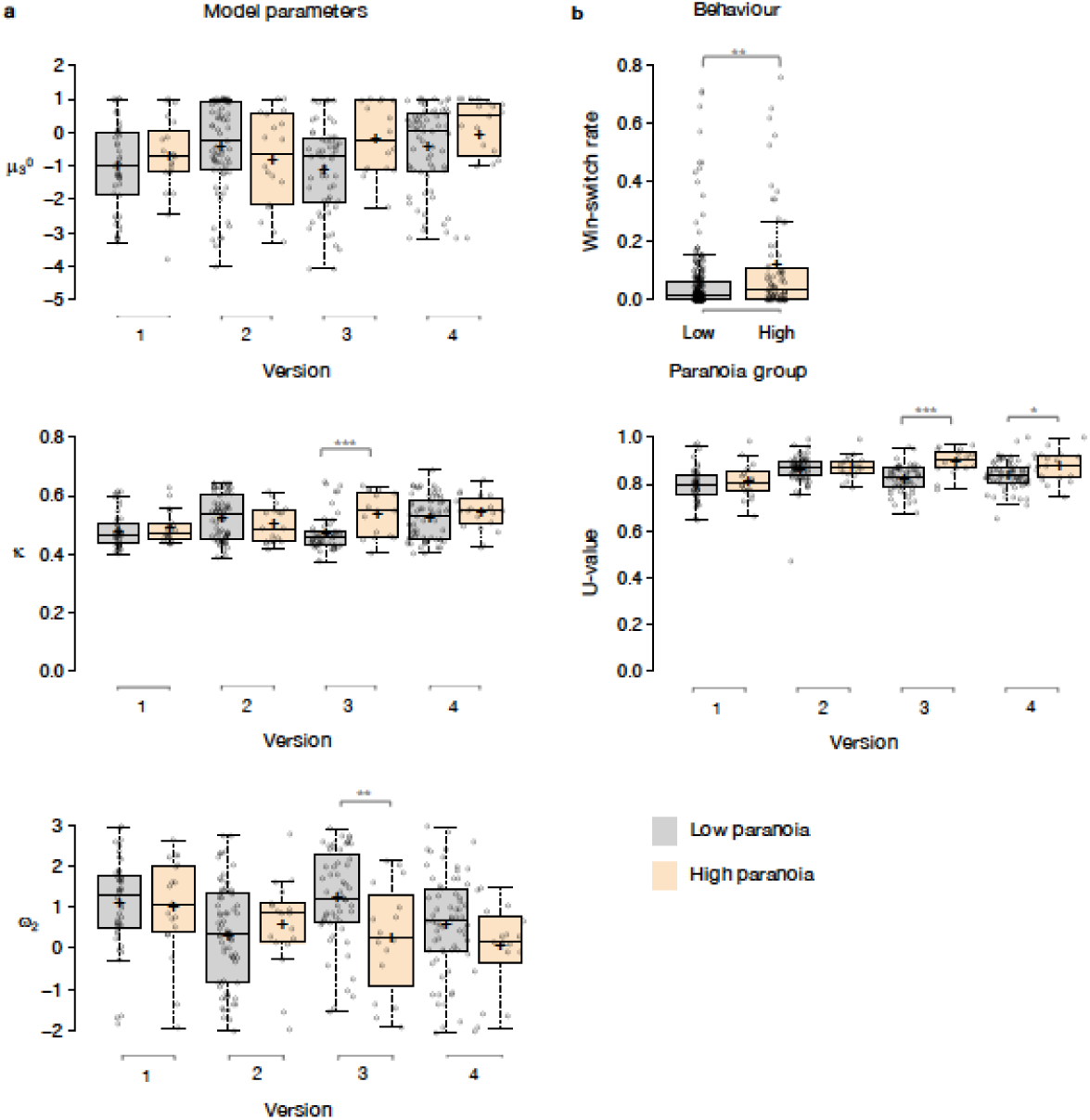
Paranoia effects across task versions. **a**, HGF parameters μ_3_^0^, κ, and ω_2_ show version 3 specific trends and effects of paranoia group membership (Experiment 2, n=234 low paranoia, 73 high paranoia, collapsed across task versions). **b**, Behaviourally, paranoid participants switched between decks more frequently after positive feedback, across all task versions and blocks (paranoia group effect; version trend, p=0.081). In versions 3 and 4 only, paranoid participants showed higher U-values, suggesting increasingly stochastic switching rather than perseverative returns to a previously rewarding option. Error bars denote standard error (SEM); *p ≤ 0.05, **p ≤ 0.01, ***p ≤ 0.001.

### Covariate analyses

We completed three ANCOVAs for each HGF parameter derived from Experiment 2: demographics (age, gender, ethnicity, and race); mental health factors (medication usage, diagnostic category, BAI score, and BDI score); and metrics and correlates of global cognitive ability (educational attainment, income, and cognitive reflection; Supplementary Tables 3 and 4). For κ, our metric of unexpected uncertainty, the paranoia by version interaction remained robust across all three ANCOVAs (demographics: *F*(3)=3.753, *P*=0.011, η_p_^2^=0.037; mental health: *F*(3)=4.417, *P*=0.005, η_p_^2^=0.049; cognitive: *F*(3)=4.304, *P*=0.005 η_p_^2^=0.043). The paranoia by version trend of μ_3_^0^ diminished with inclusion of demographic, mental health, and cognitive covariates (demographic: *F*(3)=1.997, *P*=0.119, η_p_^2^=0.020; mental health: *F*(3)=1.942, *P*=0.123, η_p_^2^=0.022; cognitive: *F*(3)=2.193, *P*=0.089, η_p_^2^=0.022). The paranoia by version interaction for ω_2_ was robust to mental health and cognitive factors (*F*(3)=3.617, *P*=0.014, η_p_^2^=0.041; *F*(3)=3.017, *P*=0.030, η_p_^2^=0.030). A paranoia group effect and paranoia by version trend remained with inclusion of demographics (ω_2_, paranoia effect: *F*(1)=4.275, *P*=0.040, η_p_^2^=0.014; interaction: *F*(3)=2.507, *P*=0.059, η_p_^2^=0.025).

### Multiple regression

We examined the effects of paranoia, anxiety, and depression on κ within the online version 3 dataset by multiple regression analysis. A significant regression equation was found (F(3,68)=3.681, p=0.016), with an R^2^ of 0.140 (Supplementary Fig.1). Participants’ predicted κ equalled 0.486 + 0.062 (PARANOIA)+0.012 (BDI) −0.006 (BAI). Paranoia was a significant predictor of κ (β=0.343, t=2.470, p=0.016, CI=[0.012, 0.113]) but depression and anxiety were not (BDI: β=0.086, t=0.423, p=0.674, CI=[-0.043, 0.066]; BAI: β=-0.043, t=-0.218, p=0.828, CI=[-0.063, 0.050]). Examination of correlation plots for κ (Supplementary Fig. 2) revealed a much stronger relationship when analyses were restricted to individuals with paranoia scores greater than 0 (i.e., endorsement of at least one item); among participants who denied all questionnaire items, a minority (seven out of 33) exhibited elevated κ. To account for the possibility that some individuals with severe paranoia may avoid disclosing sensitive information, we performed additional analyses of participants who endorsed one or more paranoia item. The correlation between paranoia and κ in the first block of the task increases from r=0.3, p=0.011, CI=[0.074, 0.497] (all participants, n=72) to r=0.588, p=8.0E-5, CI=[0.335, 0.762] (participants with paranoia > 0, n=39). In this subset, a significant regression equation was also found (F(3,35)=6.322, p=0.002), with an R^2^ of 0.351 (Supplementary Fig.1). Participants’ predicted κ was equal to 0.432 + 0.150 (PARANOIA)+0.013 (BDI) −0.004 (BAI). Paranoia was a significant predictor of κ (β=0.538, t=2.983, p=0.005, CI=[0.048, 0.252]) but depression and anxiety were not (BDI: β=0.111, t=0.494, p=0.624, CI=[-0.041, 0.067]; BAI: β=-0.035, t=-0.163, p=0.872, CI=[-0.057, 0.049]).

### Behaviour and simulations

Win-switching was the prominent behavioural feature of both paranoid participants and rats exposed to methamphetamine (Table 1, Table 2,^32^). Collapsed across blocks and task versions, our Experiment 2 data demonstrated a main effect of paranoia group (Fig. 3b; *F*(1)=9.207, *P*=0.003, η_p_^2^=0.030, MD=0.059, CI=[0.021, 0.097]; version trend: *F*(3)=2.263 *P*=0.081, η_p_^2^=0.022; low paranoia: *m=*0.06 [0.01], high paranoia: *m=*0.12 [0.02]). To elucidate whether this behaviour was stochastic or predictable (e.g., switching back to a previously rewarding option), we calculated U-values^39^, a metric of behavioural variability employed by behavioural ecologists (increasingly an inspiration for human behavioural analysis^40^), particularly with regards to predator-prey relationships^41^. When a predator is approaching a prey animal, the prey’s best course of action is to behave randomly, or in a *protean* fashion, in order to evade capture^41^. The more protean or stochastic the behaviour, the closer to the U-value is to 1. Across task blocks, paranoid participants exhibited elevated choice stochasticity (paranoia by version interaction, *F*(3)=3.438, *P*=0.017, η_p_^2^=0.033; Table 2). Post-hoc tests indicate that this stochasticity was specific to versions with contingency context change, suggesting a relationship to unexpected uncertainty (Fig. 3b; version 3, *F*(1)=17.585, *P*=3.6E-5, η_p_^2^=0.056, MD=0.071, CI=[0.038, 0.104]; version 4, *F*(1)=6.397, *P*=0.012, η_p_^2^=0.021, MD=0.039, CI=[0.009, 0.07]).

To test the propriety of our model, we simulated data for each subject in online version 3 and determined whether or not key behavioural effects (Fig. 4a, Table 1, Supplementary Table 5) were present. Using individually estimated HGF parameters to generate ten simulations per participant, we recapitulated both elevated win-switch behaviour (paranoia effect, *F*(1)=15.394, *P*=2.01E-4, η_p_^2^=0.180, MD=0.186, CI=[0.091, 0.28]) and choice stochasticity (U-value; paranoia effect, F(1)=13.362, *P*=0.0005, η_p_^2^=0.160, MD=0.065, CI=[0.030, 0.101]) in simulated paranoid participants (Fig. 4b; simulated win-switch rate, low paranoia: *m*=0.24 [0.02], high paranoia: *m*=0.43 [0.04]; simulated U-value, low paranoia: *m*=0.851 [0.008], high paranoia: *m*=0.916 [0.016]). Neither real nor simulated data showed any significant relationship between lose-stay behaviour and paranoia (Table 1, Table 2, Supplementary Table 5). To demonstrate the effects of parameters on task performance, we performed additional simulations in which we doubled or halved a single parameter at a time from the baseline average of low paranoia participants. These results confirmed the impact of κ, ω_2_, and ω_3_ on win-shift behaviour (Supplementary Fig. 3). Parameter recovery revealed significant correlations for κ and ω_2_ between original subject parameters and those estimated from simulations (Supplementary Fig. 4; ω: r=0.702, p=2.52E-11, CI=[0.557, 0.805]; κ: r=0.305, p=0.011, CI=[0.072, 0.506]). Higher level parameters (ω_3_, μ_3_^0^) were less consistently recovered, as noted in previous publications^42^.

**Fig. 4.**
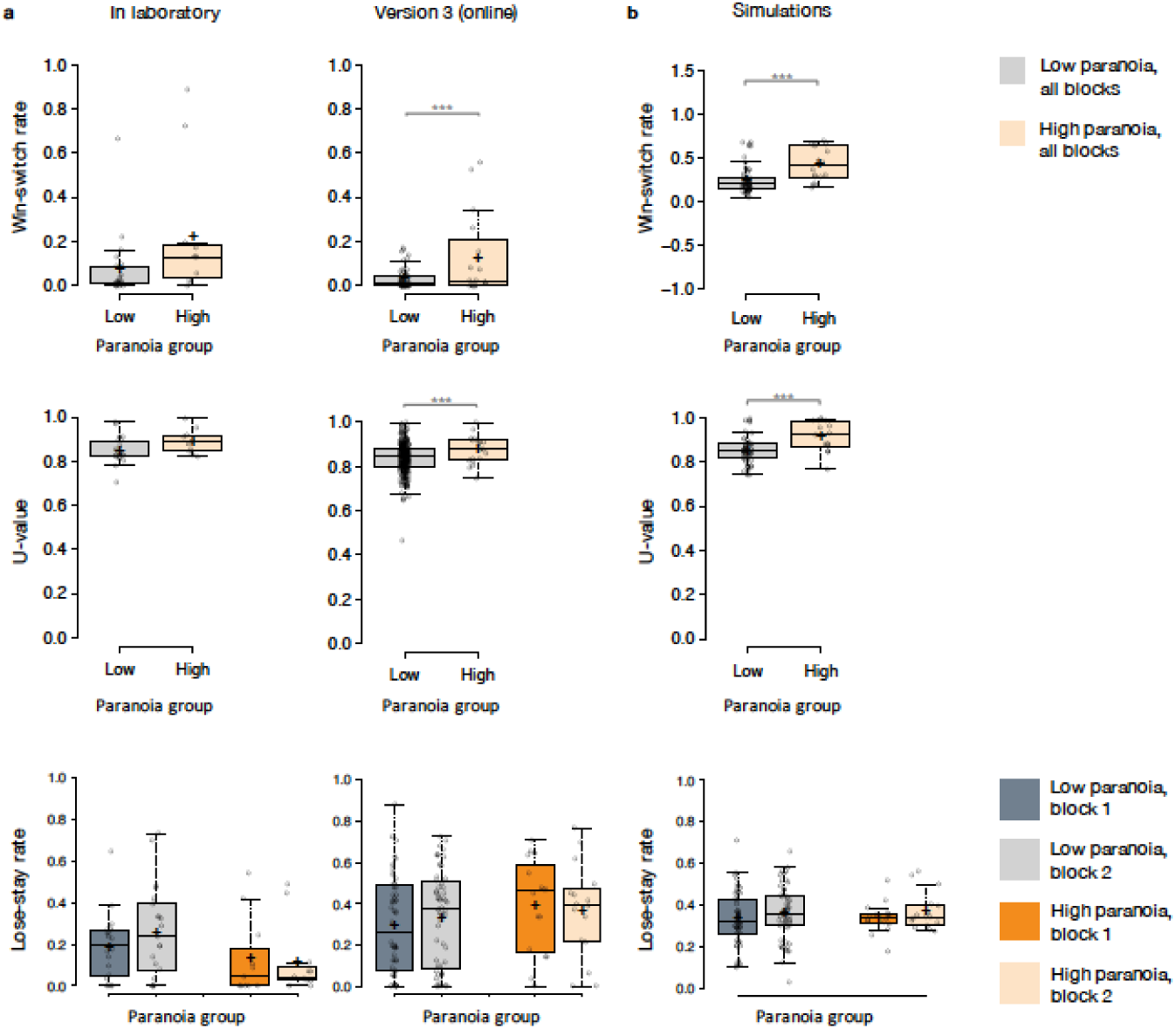
Behavioural data and simulations. **a**, Behavioural switching patterns replicate across in laboratory and online version 3 experiments (win-switch: in laboratory paranoia group trend, p=0.068; version 3 paranoia effect; U-value: in laboratory paranoia group trend, p=0.079; version 3 paranoia effect). Perseveration after negative feedback (lose-stay behaviour) did not significantly differ between paranoia groups or task block. **b**, Simulated data generated from HGF perceptual parameters (version 3) replicates win-switch and U-value behaviours (win-switch paranoia effect; U-value paranoia effect). Ten simulations were performed per subject. Rates and U-values were averaged across simulations. Error bars denote standard error (SEM); n=21 low paranoia, 11 high paranoia (in laboratory); n=56 low paranoia, 16 high paranoia (online, version 3); *p ≤ 0.05, **p ≤ 0.01, ***p ≤ 0.001.

### Clustering analysis

Given the apparent similarity in effects of paranoia and methamphetamine in humans and rats, respectively (Fig. 2b), we formally tested for latent structure in our data using two-step cluster analysis^43^. This approach automatically determines the optimal number of clusters. We analysed μ_3_^0^, κ, ω_2_, and ω_3_ estimates derived from the first block of experiment 1 and online version 3 (pre-context change data, because rats do not experience a context shift) with Post-Rx rat data. We identified two clusters with good cohesion and separation (average silhouette coefficient=0.7; cluster size ratio=2.46; Fig. 5a). All parameters contributed to clustering; κ contributed most strongly (Fig. 5b). Relative to the overall distribution, Cluster 1 was characterized by high κ and μ_3_^0^, and decreased ω_2_ and ω_3_. Cluster 2 parameters fell close to the overall distribution median, with κ and μ_3_^0^ scores lower than Cluster 1, ω_2_ and ω_3_ higher (Fig. 5a). Cluster 1 membership was significantly associated with high paranoia and methamphetamine exposure, χ^2^(1, *n=*121)=29.447, *P*=5.75E-8, Cramer’s V=0.493 (Fig. 5c). Notably, no participants in the low paranoia group with paranoia scores above zero were ascribed Cluster 1 membership. The cluster solution was robust to validation by split-half analysis, removal of the rat subjects, and removal of human participants (Supplementary Fig. 5).

**Fig. 5.**
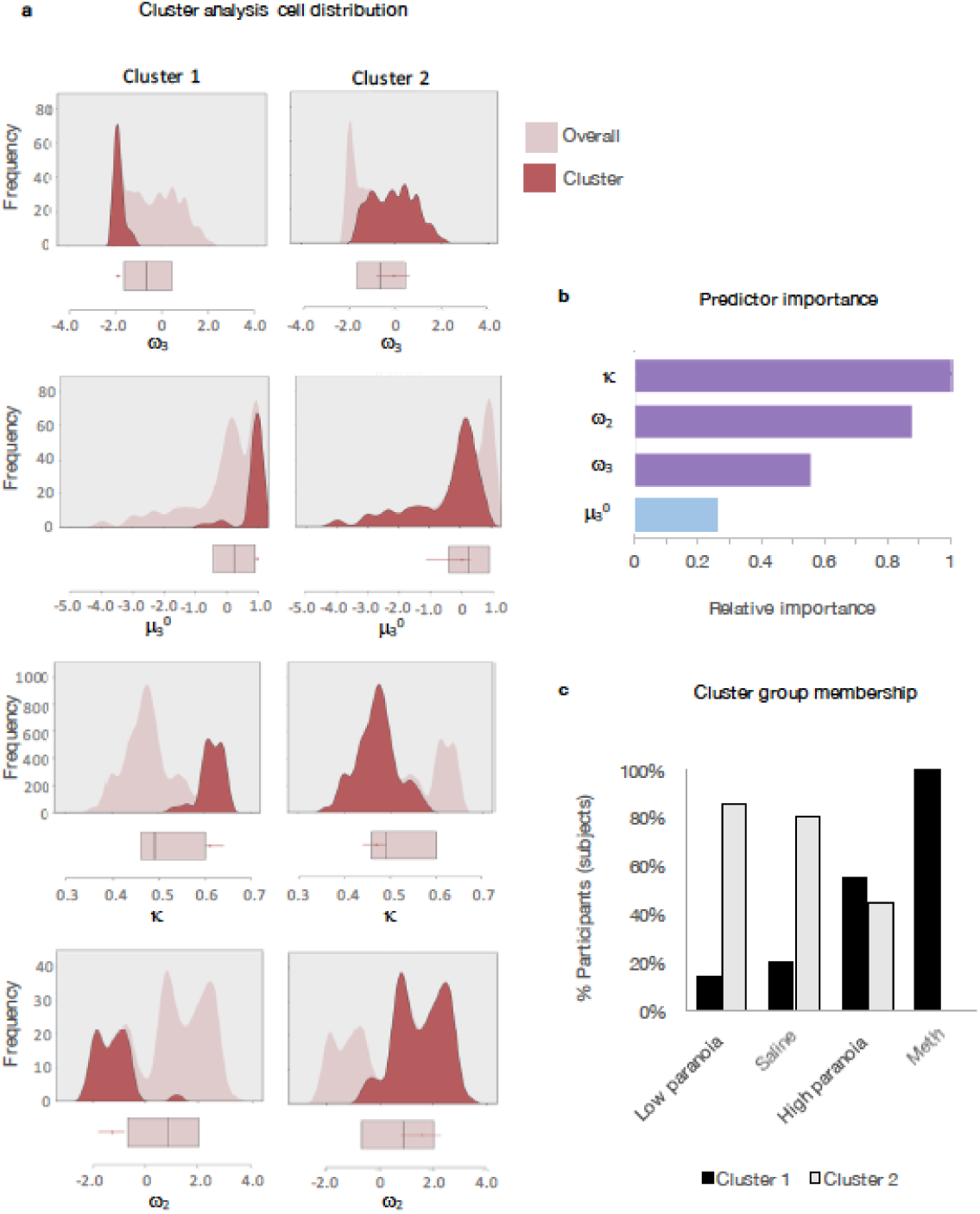
Cluster analysis of HGF parameters. Two-step cluster analysis of model parameters across rat and human data sets (rat, post-Rx; in laboratory and online version 3, block 1). Automated clustering yielded an optimal two clusters with good cohesion and separation (average silhouette coefficient=0.7; cluster size ratio=2.46). **a**, Parameter density plots for overall distributions (light pink) and cluster-specific distributions (red). Box-plots of overall median, 25^th^ quartile, and 75^th^ quartile are aligned below each plot (pink), with cluster medians and quartiles superimposed (red). Relative to the overall distribution, Cluster 1 (n=35) medians are elevated for μ_3_^0^ and κ, decreased for ω_2_ and ω_3_. Cluster 2 (n=86) falls within each overall distribution. **b**, Predictor importance of included parameters. **c**, Distribution of cluster identities within groups (low paranoia: n=77; high paranoia: n=27; rat-saline: n=10; rat-methamphetamine: n=7). Cluster 1 membership is significantly associated with paranoia and methamphetamine groups (χ^2^(1, n=121)=29.447, p=5.75E-8).

## Discussion

We have shown that inferential learning differs in paranoid individuals according to a specific pattern of beliefs about contextual volatility and response to uncertainty. During probabilistic reversal-learning with three options, paranoid individuals and rats chronically exposed to methamphetamine have higher initial expectations of task volatility (μ_3_^0^). In other words, they start the task anticipating more changes in stimulus-outcome associations. These same subjects respond more strongly to perceived volatility in updating stimulus-outcome associations (κ). This coincides with decreased ω_2_, reflecting more stable beliefs about the underlying probability values themselves. Consequently, inferred contextual volatility manifests in excessive perception of reversal events, increasing behavioural switching.

We replicated an elevated prior on environmental volatility (μ_3_^0^) and higher sensitivity to this volatility (κ) previously observed in HGF analyses of 2-choice probabilistic reversal-learning in medicated and unmedicated patients with schizophrenia^44^. Unlike prior work, we assessed the trans-diagnostic symptom of paranoia across the continuum of health and illness, provided three choice options to differentiate stochastic switching from perseverative returns, and explicitly manipulated unexpected uncertainty across task versions. The version that shifts from an easier to discern contingency context to a more difficult context was associated with paranoia group differences in μ_3_^0^, κ, and ω_2_. Furthermore, this context change elicited decreases in metavolatility learning (ω_3_) among low paranoia controls relative to their first block baseline, rendering them more similar to high paranoia participants. Paranoid individuals behave as if that the world is always more volatile, demanding continual updating of associations. Low paranoia individuals behave similarly under more difficult, uncertain conditions. Although our domain-general paradigm lacks any sizable, tangible threat, uncertainty itself may be aversive^45^, threatening the brain’s ability to make clear predictions about future states and actions.

Unexpected uncertainty, the perception of change in the probabilities of the environment — particularly “unsignaled context switches”^25^ — is thought to promote abandonment of old associations and new learning. Our analysis of covariates warrants specific focus on κ, the sensitivity to unexpected uncertainty. Other parameter-paranoia associations did not endure after controlling for demographic factors (age, gender, ethnicity, and race). These factors are strong predictors of paranoia^46-48^. It is notable too that κ was the most powerful discriminator of the two clusters of human and animal participants. However, the rodent data are less impacted by concerns about covariates, and there, the other parameters (prior on volatility, expected uncertainty, meta-volatility) were all changed by methamphetamine. We conclude that κ is the parameter most robustly associated with paranoia.

Multiple neurobiological manipulations may induce win-switching behaviour. Lesions of the mediodorsal thalamus in non-human primates^49^ or neurons projecting from the amygdala to orbitofrontal cortex in rats^50^ engender win-switching. The human hippocampus appears to be sensitive to volatility during belief learning, as are the anterior cingulate cortex and insula^27^. However, unexpected uncertainty, and the κ parameter of the HGF in particular^51^, are thought to be signalled via the locus coeruleus and noradrenaline (i.e., neural gain)^25-28^. This mechanism is thought to coordinate rapid shifts in cortical networks through patterns of widespread norepinephrine release, modulating exploratory versus exploitative behaviours (i.e., switching and staying)^52-55^ and responding to stress^56-58^, unexpected uncertainty^25^,^27^ and subliminal fear cues^59^ to coordinate fight-or-flight responses^58^. In fact, visual fear stimuli presented below the threshold of conscious perception activate the locus coeruleus, amygdala, and fronto-temporal orienting regions, suggesting a neural ‘alarm’ system for rapid threat detection^59^. The dual role of the locus coeruleus in recognizing and responding to threats as well as unexpected uncertainty suggests that dysfunction could produce both paranoia and the inferential abnormalities we observed. Methamphetamine may induce similar dysfunction. Acute moderate doses increase pre-synaptic catecholamine release, particularly noradrenaline^60^, and induce exploratory locomotive effects modulated through adrenoceptors on dopamine neurons^61^. Unlike acute binge paradigms, the schedule of methamphetamine administration completed by the rats in these analyses preserves methamphetamine-induced locomotor hyperactivity^32,62^.

Perturbations of noradrenergic gain impede new learning while appearing falsely to enhance behavioural flexibility. In rats, excessive release of noradrenaline from the locus coeruleus into the anterior cingulate cortex drives disengagement from model-consistent performance in a three-option counter prediction task^28^. This manipulation includes stochastic switching and insensitivity to feedback or context change— a type of behavioural “flexibility” that is ultimately inflexible. Our data suggest that in paranoia, increased gain under uncertainty may similarly shunt away incoming information, leaving only reflexive, habitual responses. Although participants engage in choice switching— in an increasingly stochastic fashion— our cluster analyses show that excessive κ is associated with diminished metavolatility learning, rendering these subjects less flexible in updating context beliefs. In this fashion, excessive switching behaviour may be indicative of fixed higher-level beliefs.

Disengagement from model-congruent behaviour has been observed in paranoia and psychosis^63,64^. Evolutionarily, departure from predictable, rational modes of behaviour might offer an adaptive mechanism for escape from intractable threat. As a protean defence mechanism, behavioural stochasticity impedes predators’ abilities to create accurate, actionable countermeasures^41,65,66^. If driven by excessive noradrenergic gain, protean defence may represent an extreme state along a heavily conserved, continuous common mechanism underlying vigilance and false alarms^67-69^, arousal-linked attentional biases and selective processing (i.e., focusing on narrow, most salient features versus broader context^70^; attending to and learning from predisposition-conforming features^55^), and behavioural and cognitive flexibility in response to unexpected uncertainty and Bayesian surprise (i.e., prediction error)^53,54,71^. We hypothesize that individuals with stable, trait-level paranoia, rather than having specific deficits in inferring others’ reputations^16^, exhibit disturbances across the domains of behavioural flexibility and stochasticity, false alarms and attentional bias, and inferential response to unexpected uncertainty. Our data suggest that that these perturbations exist outside of social settings and may be elicited in nonhuman models. We propose that protean defence and its attendant behavioural stochasticity might be one useful translational marker of paranoia.

We conclude that this model provides a robust tool for computational dissection of learning mechanisms across species. Social interactions play a rich and undeniable role, but translational, domain-general approaches may ultimately facilitate biological insights into paranoia, psychosis and delusions^72,73^. Whilst we contend that our task is relatively free of social features (certainly compared to others^15^), the possibility remains that the elevated U-values in our participants are reflective of attempts (and perhaps failures) to predict our intentions as experimenters. Indeed, this is a possibility raised previously with regards to simple conditioned behaviours in experimental animals. Even during Pavlovian conditioning, animals may attempt to infer a generative model of the task environment, which might, ultimately, include the experimenter arranging the contingencies^74,75^. It is possible that all instances of human cognitive testing involve an element of inference by the participant with regards to the intentions of the experimenter, whether or not the task at hand is explicitly social, and indeed, all cognitive functions may be aimed at or modulated by such inferences^76^.

In summary, a strong belief in the volatility of the world necessitates hypervigilance and a facility with change. However, in paranoia, that belief (in the volatility of the world) is itself resistant to change, making it difficult to reassure, teach, or change the minds of people who are paranoid. They remain “on guard” even under stable conditions. Whether promoting recovery from paranoia-associated illness^77^ or interpersonal collaboration, our domain-general approach reaffirms the merit of trying to establish stable, predictable environments. We note with interest the apparent relationship between conspiratorial ideation and societal crisis situations (terrorist attacks, plane crashes, natural disasters or war) throughout history, with peaks around the great fire of Rome (AD 64), the industrial revolution, the beginning of the cold war, 9/11, and contemporary financial crises^78^. Perhaps these broader trends are a macrocosmic version of the unexpected uncertainty manipulation that drove promiscuous switching behaviour in our task, particularly in high paranoia participants. Rather than proving adaptive, their behaviour ultimately increases the noise of their task experience with sampling of sub-optimal options and exposure to misleading positive feedback. In today’s world of escalating uncertainty and volatilty – particularly environmental climate change – our findings suggest that the paranoid style of inference may prove particularly maladaptive for coordinating collaboratve solutions.

## Methods

Experiments were conducted at Yale University and the Connecticut Mental Health Center (New Haven, CT) in strict accordance with Yale University’s Human Investigation Committee and Institutional Animal Care and Use Committee. Informed consent was provided by all research participants.

**Experiment 1**. English-speaking participants aged 18 to 65 (*n=*34) were recruited from the greater New Haven area through public fliers and mental health provider referrals. Exclusion criteria included history of cognitive or neurologic disorder (e.g., dementia), intellectual impairment, or epilepsy; current substance dependence or intoxication; cognition-impairing medications or doses (e.g. opiates, high dose benzodiazepines); history of special education; and colour blindness. Participants were classified as healthy controls (*n=*18), schizophrenia spectrum patients (schizophrenia or schizoaffective disorder; *n=*8), and mood disorder patients (depression, bipolar disorder, generalized anxiety disorder, post-traumatic stress disorder; *n=*8) on the basis of clinician referrals and/or self-report. Participants were compensated $10 for enrolment with an additional $10 upon completion. Two healthy controls were excluded from analyses due to failure to complete the questionnaires and suspected substance use, respectively.

**Experiment 2**. 332 participants were recruited online via Amazon Mechanical Turk (MTurk). The study advertisement was accessible to MTurk workers with a 90% or higher HIT approval rate located within the United States. To discourage bot submissions and verify human participation, we required participants to answer open-ended free response questions; submit unique, separate completion codes for the behavioural task and questionnaires; and enter MTurk IDs into specific boxes within the questionnaires. All submissions were reviewed for completion code accuracy, completeness of responses (i.e., declining no more than 30% of questionnaire items), quality of free response items (e.g., length, appropriate grammar and content), and use of virtual private servers (VPS) to submit multiple responses and/or conceal non-US locations (Dennis VPS paper, 2018). Upon approval, workers were compensated $6. Those who scored in the top 25% on the card game (reversal-learning task) earned a $2 bonus. We rejected or excluded 19 submissions that geolocation services (https://www.iplocation.net/) identified as originating outside of the United States or from suspected server farms, 4 submissions for failure to manually enter MTurk ID codes, and 2 submissions for insufficient questionnaire completion. Submissions with grossly incorrect completion codes were rejected without further review.

**Experiment 3.** Subject information, behavioural data acquisition, and behavioural analyses were described previously ^32^. Long Evans rats (Charles River; *n=*20) ranged from 7 to 9 weeks of age. Rats were exposed to escalating doses and frequency of saline (*n=*10) or methamphetamine (*n=*10, 3 withdrawn during dosing), imitating patterns of human methamphetamine users^62,79^. Prior to dosing (Pre-Rx), rats completed 26 within-session reversal sessions, including up to 8 reversals per session. Post-dosing (Post-Rx), rats completed one test session per week for four weeks. Computational model parameters were estimated from each session and averaged across treatment conditions to yield one Pre-Rx and Post-Rx set of parameters per rat.

### Behavioural task

Participants completed a 3-option probabilistic reversal-learning paradigm. Three decks of cards were displayed on a computer monitor for 160 trials. Participants selected a deck on each trial by pressing the predesignated key. We advised participants that each deck contained winning and losing cards (+100 and −50 points), but in different amounts. We also stated that the best deck may change. Participants were instructed to find the best deck and earn as many points as possible. Probabilities switched between decks when the highest probability deck was selected in 9 out of 10 consecutive trials (performance-dependent reversal). Every 40 trials the participant was provided a break, following which probabilities automatically reassigned (performance-independent reversal).

In Experiment 1, the task was presented via Eprime® 2.0 software (Psychology Software Tools, Sharpsburg, PA). Participants were limited to a 3-second response window, after which the trial would time out and record a null response. A fixation cross appeared during variable inter-trial intervals (jittering). Task pacing remained independent of response time. In block 1 (trials 1-80) the reward probabilities (contingency context) of the three decks were 90%, 50%, and 10% (90-50-10%). Without cue or warning, the context changed to 80%, 40%, and 20% (80-40-20%) upon initiation of block 2 (trials 81-160).

In Experiment 2, the task was administered via web browser link from the MTurk marketplace. We changed the task timing to self-paced and eliminated null trials and inter-trial jittering. A progress tracker was provided every 40 trials. Workers were randomly assigned to one of four task versions. Version 1 had a constant contingency context of 90-50-10%. Version 4 maintained a constant context of 80-40-2. Version 3 replicated the 90-50-10% (block 1) to 80-40-20% (block 2) context change of Experiment 1. Version 4 presented the reversed context change, 80-40-20% (block 1) to 90-50-10% (block 2). We analysed attrition rates across the four versions.

### Questionnaires

Following task completion, questionnaires were administered via the Qualtrics® survey platform (Qualtrics Labs, Inc., Provo, UT). Items included demographic information (age, gender, educational attainment, ethnicity, and race) and mental health questions (past or present diagnosis, medication use, *Structured Clinical Interview for DSM-IV Axis II Personality Disorders* (SCID-II)^33^, Beck’s Anxiety Inventory (BAI)^34^, Beck’s Depression Inventory (BDI)^35^. We removed the single suicidality question from the BDI for Experiment 2. Experiment 2 included additional items: income, three cognitive reflection questions (Supplementary Table 4), and three free response items (‘What do you think the card game was testing?’, ‘Did you use any particular strategy or strategies? If yes, please describe’, and ‘Did you find yourself switching strategies over the course of the game?’). We quantified trait-level paranoia using the paranoid personality subscale of the SCID-II, and we included an ideas of reference item from the schizotypy subscale (‘When you are out in public and see people talking, do you often feel that they are talking about you?’) This item, along with other SCID-II items, has previously been included as a metric of paranoia in the general population^5,80^. Participants who endorsed 4 or more paranoid personality items (i.e., the cut-off for the top third identified in Experiment 1) were classified as ‘high paranoia.’ Each participant’s SCID-II, BAI, and BDI scores were normalized by total scale items answered. Distributions of SCID-II scores are shown in Supplementary Fig. 6. Response rates were higher than 90% for all questionnaire items and scales (Supplementary Table 6).

### Behavioural analysis

We analysed tendencies to choose alternative decks after positive feedback (win-switch) and select the same deck after negative feedback (lose-stay). Win-switch rates were calculated as the number of trials in which the participant switched after positive feedback divided by the number of trials in which they received positive feedback. Lose-stay rates were calculated as number of trials in which a participant persisted after negative feedback divided by total negative feedback trials. In Experiment 1, we excluded post-null trials from these analyses. To further characterize switching behaviour, we calculated U-values, a measure of choice stochasticity:

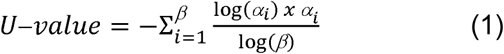

where *β* is the number of possible choice options (i.e., card decks or noseports) and *α* equals the relative frequency of choice option *i* ^39^. To avoid any choice counterbalancing effects across reversals, choice frequencies were determined by the underlying probabilities of the decks rather than their physical attributes (e.g., deck position or colour). Additional behavioural analyses included trials to first reversal, trials to post-reversal recovery, and trials to post-reversal switch. The latter two were restricted to the first reversal in the first block. Trials post-reversal were counted from the first-negative feedback trial following the true reversal event. Recovery was defined as switching to the best deck and staying for at least one additional trial.

### Computational modelling and simulations

We utilized the freely available HGF toolbox v5.3.1 (https://translationalneuromodeling.github.io/tapas/) in MATLAB and Statistics Toolbox Release 2016a (MathWorks ®, Natick, MA)^30,31^. The HGF employs an “observing the observer” framework consisting of a perceptual model (a generative model of the agent’s inferences about the task environment) and a decision model that reconciles the agent’s actions. HGF parameter values are inferred from observed agent decisions and trial-by-trial feedback (i.e., win or loss outcomes) through variational model inversion^30,31^. Our model schema consisted of a 3-level HGF multi-arm bandit configuration for binary outcomes, paired with the softmax-mu03 decision model. The softmax mu03 model tests the hypothesis that beliefs about environmental volatility dynamically influence behaviour. The inverse decision temperature (β) is set to the inverse volatility estimate exp (-μ_3_^(*k*)^) where *k* denotes the current trial, permitting simultaneous estimation of μ^0^, κ and ω. We inspected each subject’s x_1_, x_2_, and x_3_ trajectories and optimized the default perceptual model configuration file by changing the second level κ prior mean from log(1) to log(0.6). The first level κ remained fixed at log(1). Third level trajectory were regularized by use of the autoregressive HGF configuration option.

Perceptual parameters were estimated separately for blocks 1 and 2, with block 1 μ_2_^0^ and μ_3_^0^ comprising the μ_2_ and μ_3_ prior means in block 2. To evaluate the validity of our model, we subsequently simulated participant choices using trial-by-trial outcome data and estimated perceptual parameters from online version 3 participants. We performed ten simulations per subject and calculated win-shift rates, U-values, and lose-stay rates to compare with our actual data. Code for parameter estimation and simulations are detailed in the Supplementary Methods.

### Statistics

Unless otherwise specified, statistical analyses and effect size calculations were performed in IBM SPSS Statistics, Version 25 (IBM Corp., Armonk, NY), with an alpha of 0.05. Box-plots were created with the web tool BoxPlotR^81^. Model parameters were corrected for multiple comparisons using the Benjamini Hochberg (False Discovery Rate) method. Bonferroni corrections were largely consistent (Supplementary Table 2)

To compare questionnaire item means between two groups (Table 1, low versus high paranoia), we conducted independent samples t-tests. To compare questionnaire item means across paranoia groups and task versions (Table 2, fixed factors), we employed univariate analyses. Associations between characteristic frequencies and subject group or task version were evaluated by Chi-Square Exact tests (two groups) or Monte Carlo tests (more than 2 groups). Pearson correlations established the associations between paranoia and BDI scores, BAI scores, win-switch rates, and κ. We selected two-tailed p-values where applicable and assumed normality. Multiple regressions were conducted with κ estimates from the first task block (dependent variable) and paranoia, BAI, and BDI scores from online version 3.

To compare HGF parameter estimates and behavioural patterns (win-switch, U-value, lose-stay) across block, paranoia group (Experiment 1, Experiment 2 version 3), and/or task version (Experiment 2), we employed repeated measures and split-plot ANOVAs (i.e., block designated within-subject factor, paranoia group and task version as between subject). We similarly evaluated Experiment 3 parameter estimates for treatment by time interactions. For Experiment 2, we performed ANCOVAs for μ_3_^0^, κ, ω_2_, and ω_3_ to evaluate three sets of covariates: (1) demographics (age, gender, ethnicity, and race); (2) mental health factors (medication usage, diagnostic category, BAI score, and BDI score); (3) and metrics and correlates of global cognitive function (educational attainment, income, and cognitive reflection). Unless otherwise stated, post-hoc tests were conducted as least significant difference (LSD)-corrected estimated marginal means.

Meta-analyses were conducted using random effects models with the R Metafor package^82^. Mean differences were assessed for low versus high paranoia groups in the in laboratory experiment and online version 3. Standardized mean differences (methamphetamine or high paranoia versus saline or low paranoia) were employed to account for the differences in task design between animal and human studies.

The 2-step clustering analysis approach was selected to automatically determine optimal cluster count and cluster group assignment. Clustering variables included paranoia-relevant parameter estimates (μ_3_^0^, κ, ω_2_, and ω_3_) from Experiment 1 (block 1); online, version 3 (block 1), and rats (Post-Rx) as continuous variables with a Log-likelihood distance measure, maximum cluster count of 15, and Schwarz’s Bayesian Criterion (BIC) clustering criterion. We validated our clustering solution by sorting the data into two halves and running separate cluster analyses. We also compared cluster solutions derived exclusively from rat data versus human data. A Chi-Square test determined the significance of the association between cluster membership and group (methamphetamine/high paranoia versus saline/low paranoia).

## Data availability

Data are available on ModelDB^83^ (http://modeldb.yale.edu/258631) with accession code **p2c8q74m**. Figures 2, 3, 4, and 5 and Supplemental Figures 1 and 2 have associated raw data.

## Code availability

Code for the HGF toolbox v5.3.1 is freely available at https://translationalneuromodeling.github.io/tapas/. Additional instructions are provided in the Supplementary Information. Task code is available by request.

## Acknowledgements

This work was supported by the Yale University Department of Psychiatry, the Connecticut Mental Health Center (CMHC) and Connecticut State Department of Mental Health and Addiction Services (DMHAS). It was funded by an IMHRO / Janssen Rising Star Translational Research Award, an Interacting Minds Center (Aarhus) Pilot Project Award, and NIMH R01MH12887 (P.R.C.). E.J.R. was supported by the NIH Medical Scientist Training Program Training Grant, GM007205; NINDS Neurobiology of Cortical Systems Grant, T32 NS007224; and a Gustavus and Louise Pfeiffer Research Foundation Fellowship. S.U. received funding from NSF Fellowships DGE1122492 and DGE1752134. S.M.G. and J.R.T. were supported by NIDA DA DA041480. The funders had no role in study design, data collection and analysis, decision to publish or preparation of the manuscript. The authors thank Dr. James Waltz for providing an earlier version of the reversal-learning e-prime code. The authors acknowledge the help, support, and advice of Dr. Sarah Fineberg, Dr. Albert Powers III, and Dr. Pantelis Leptourgos.

## Author contributions

E.J.R., S.U., C.D.M., J.R.T., S.M.G., and P.R.C. contributed to the conception and design of the experiment. S.U. and E.J.R. developed the online experiments. C.D.M. advised on modelling analyses. S.M.G. and J.R.T. contributed the rodent dataset. E.J.R. conducted the experiments, collected the data, and analysed the data. E.J.R. and P.R.C. drafted the manuscript. All authors reviewed the manuscript and gave final approval for publication.

---

**Supplementary Fig. 1.**
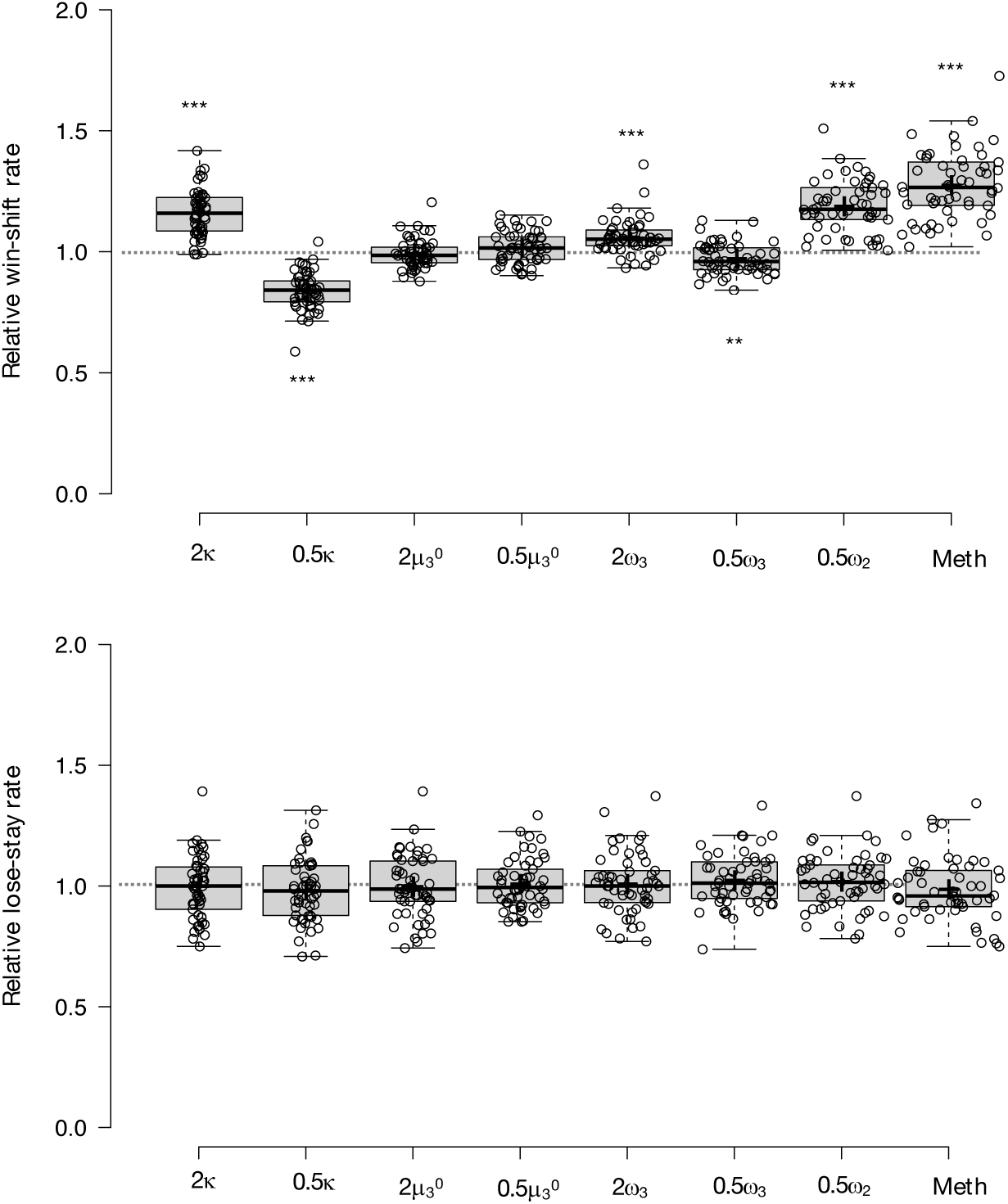
Parameter effects on simulated task performance. We simulated behaviour from low paranoia participants (online Version 3, n=54) to evaluate the effects of **κ**, **μ**_3_^0^, **ω**_2,_ and **ω**_3_ on win-shift and lose-stay rates. Estimated perceptual parameters were averaged across subjects to create a single set of baseline parameters. Additional parameter sets were created by doubling or halving one parameter at a time (e.g., 2 **κ** or 0.5 **κ**), while the others were held constant (n.b., 2 **ω**_2_ violated model assumptions and was excluded from analysis). We also included the average parameter values of rats exposed to methamphetamine (Meth). Ten simulations were run per subject for each condition (i.e., parameter set). Win-shift and lose-stay rates were calculated, then averaged across simulations and subjects. Rates from each condition were divided by the baseline condition rate to generate relative win-shift and lose-stay rates. We compared relative rates for each condition to the baseline (relative rate of 1; paired t-tests, Bonferroni-corrected p-values). Baseline parameters were positive for **κ** and **ω**_2,_ and negative for **μ**_3_^0^ and **ω**_3_. Consequently, the doubled (2x) condition makes **μ**_3_^0^ and **ω**_3_ more negative (lower). (n=54). Error bars denote standard error (SEM); *p ≤ 0.05, **p ≤ 0.01, ***p ≤ 0.001.

**Supplementary Fig. 2.**
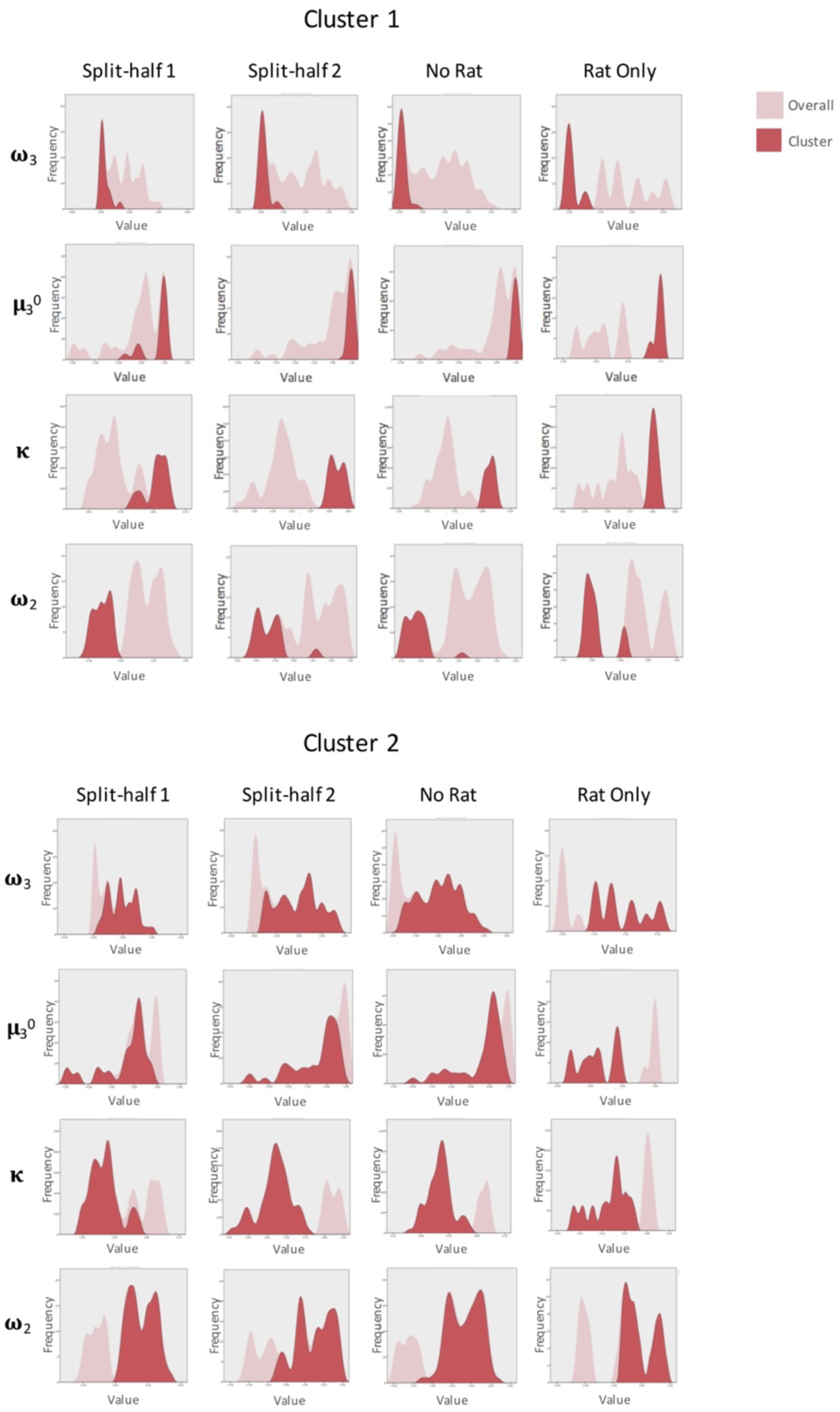
Cluster validation. We replicated our 2-cluster solution (Fig. 4) by independently running two-step cluster analyses on separate halves of the data (Split-half 1, Split-half 2), removing the rat data and running the human data only (No Rat), and running the rat data alone (Rat Only). In each condition, we identified two clusters with good cohesion and separation (Split-half 1, n=19 cluster 1, 42 cluster 2: silhouette coefficient = 0.6; Split-half 2, n = 17 cluster 1, 43 cluster 2: silhouette coefficient = 0.7; No Rat, n=26 cluster 1, 78 cluster 2: silhouette coefficient = 0.7; Rat Only, n=6 cluster 1, 11 cluster 2: silhouette coefficient = 0.7). All variables showed predictor importance above 0.2 with some variation in order of importance (Split-half 1, **ω**_2_ > **κ** > **ω**_3_ > **μ**_3_^0^; Split-half 2, **κ** > **ω**_2_ > **ω**_3_ > **μ**_3_^0^; No Rat, **κ** > **ω**_2_ > **ω**_3_ > **μ**_3_^0^; Rat Only, **μ**_3_^0^ > **ω**_2_ > **ω**_3_ > **κ**). Predictor importance was weighted more evenly across variables in the Rat Only condition; all variables showed predictor importance above

