## Supplement for "Expecting the unexpected: the paranoid style of belief updating across species"

*Philip R. Corlett

**This PDF file includes:**

Supplementary Methods

Supplementary Figures 1 through 6

Supplementary Tables 1 through 7

References for supplementary methods

**Supplementary methods**

Computational modelling

**Materials.** The Hierarchical Gaussian Filter (HGF) toolbox v5.3.1 is freely available for download in the TAPAS package at <https://translationalneuromodeling.github.io/tapas>^1,2^. We installed and ran the package in MATLAB and Statistics Toolbox Release 2016a (MathWorks ®, Natick, MA).

**Perceptual parameter estimation.** In the human reversal-learning experiments, we estimated perceptual parameters individually for the first and second halves of the task (i.e., blocks 1 and 2). Each participant’s choices (i.e., deck 1, 2, or 3) and outcomes (win or loss) were entered as separate column vectors with rows corresponding to trials. Wins were encoded as ‘1’, losses as ‘0’, and choices as ‘1’, ‘2’, or ‘3’. We selected the autoregressive 3-level HGF multi-arm bandit configuration for our perceptual model and paired it with the softmax-mu03 decision model. For each participant, block 1 parameters were estimated as follows:

>> est = tapas_fitModel(choices, outcomes, 'tapas_hgf_ar1_binary_mab_config','tapas_softmax_mu3_config')

where choices is the column vector of the participant’s choices for block 1, and outcomes is the column vector of the corresponding outcomes. We then plotted the 3-level belief trajectory:

>> tapas_hgf_ar1_binary_mab_plotTraj(est)

After checking trajectories and log model evidence scores to find perceptual parameter configurations that fit all participants, we optimized the default configuration file (‘tapas_hgf_ar1_binary_mab_config.m’) by changing line 174 to

c.logkamu = [log(1), log(0.6)];

We ran all block 1 parameter estimates with this configuration. For block 2, we implemented an additional change to the configuration file. For each subject, we set the prior mean start points for 𝛍_2_ and 𝛍_3_ (line 147, ‘c.mu_0mu = [NaN, 0, 1]’) to the values of 𝛍_2_^0^ and 𝛍_3_^0^ estimated in block 1.

Rat reversal-learning data was entered similarly, with choices designated as ‘1’, ‘2’, or ‘3’ and reward presence or absence noted as ‘1’ and ‘0’, respectively. Perceptual parameters were estimated as a single block per session and averaged across Pre-Rx or Post-Rx sessions for each subject. Since the contingency context remained 70-30-10%, we used the default start point values of 𝛍_2_ and 𝛍_3_ in ‘tapas_hgf_ar1_binary_mab_config.m’ (i.e., the same configuration file used in block 1 estimations for the human reversal-learning experiments). Parameters were estimated with the same code as described above.

**Simulations.** We performed ten simulations per participant (online version 3) to determine whether our parameter estimates and model successfully captured behavioral differences between groups (e.g., win-switch rates). Each simulation required the participant’s actual data (i.e., the column vectors ‘outcomes’ and ‘choices’) and the corresponding set of derived perceptual parameters (‘est.p_prc.p’). On each trial, a new choice was simulated conditional on the actual inputs in previous trials. The following performs a single simulation, yielding an artificial choice column vector [‘sim.y’]:

>> sim = tapas_simModel([outcomes, choices], 'tapas_hgf_ar1_binary_mab', est.p_prc.p, 'tapas_softmax_mu3', []);

We performed an additional supplementary analysis (Fig. S1) to illustrate the effects of each parameter on task behavior, i.e., by doubling or halving one parameter at a time. First we established a baseline set of perceptual parameters containing the average values from the low paranoia participants (online version 3). We then ran 10 simulations per subject for each of the following conditions: baseline, 2𝛋, 0.5𝛋, 2𝛍_3_^0^, 0.5𝛍_3_^0^, 2𝛚_3_, 0.5𝛚_3_, 2𝛚_2_, 0.5𝛚_2_, and the average perceptual parameters (𝛋, 𝛍_3_^0^, 𝛚_3_, and 𝛚_2_) from Post-Rx methamphetamine rats. The 2𝛚_2_ condition yielded parameters in a region where model assumptions were violated (negative posterior precision error message) and was excluded from further analysis. Win-shift and lose-stay rates were calculated from each simulation as follows, and then averaged for each condition:

$$Win\text{-}switch rate=\frac{Number of trials in which choice switched after positive feedback}{Total positive feedback trials}$$

$$Lose\text{-}stay rate=\frac{Number of trials in which choice repeated after negative feedback}{Total negative feedback trials}$$

For each participant, we divided rates derived from each condition by the baseline rates to determine relative win-switch and lose-stay rates. We compared each relative rates to the baseline condition (i.e., 1.0) with paired-samples t-tests using Bonferroni-corrected p-values.

Parameter recovery. We performed perceptual parameter estimation (see above) on 10 simulations per subject using first block data from online version 3. These simulations were generated from each subject’s corresponding perceptual parameters. We averaged recovered parameters across simulations and low versus high paranoia.

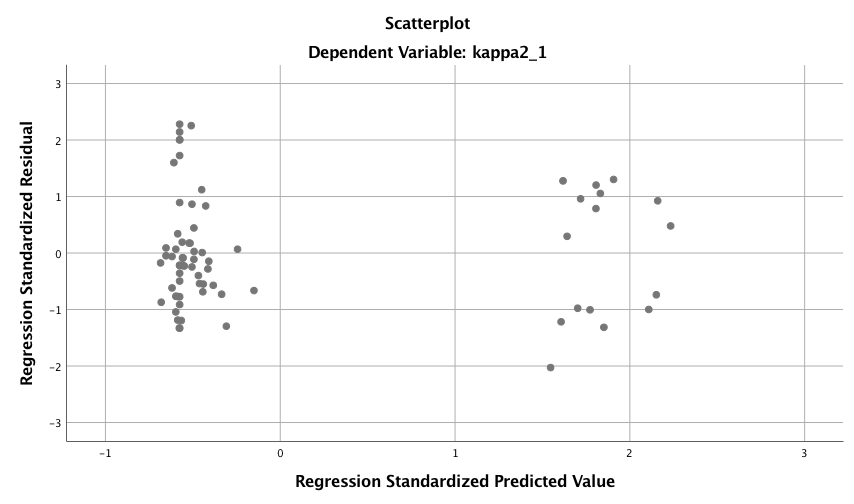

**a**

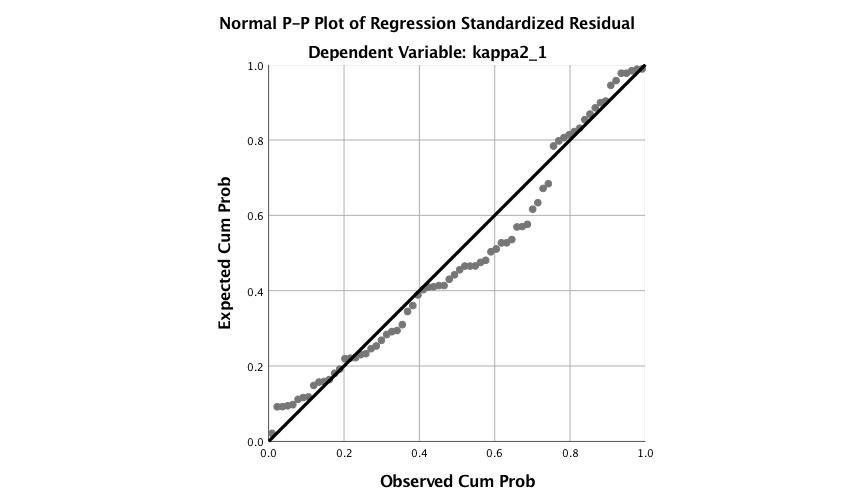

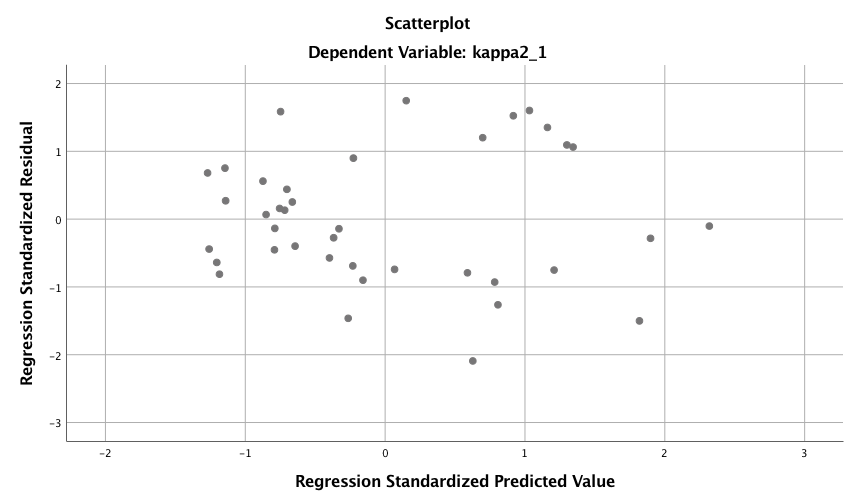

**bA**

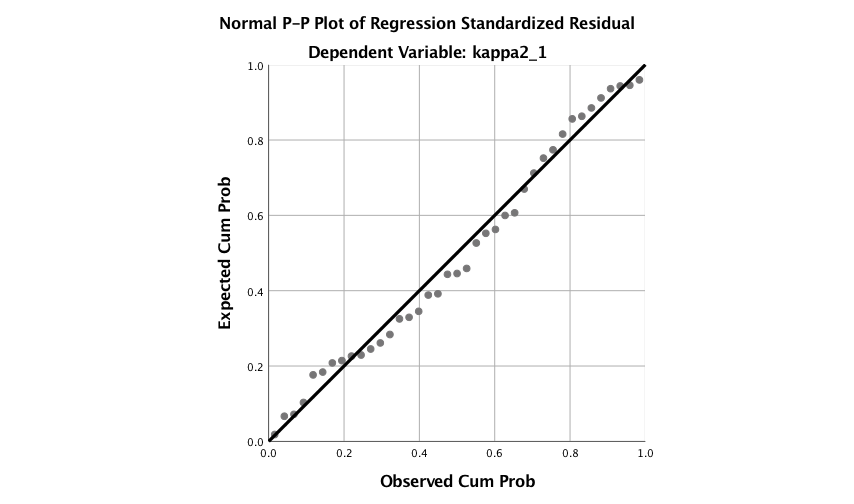

**Supplementary Fig. 1. Multiple regression analyses for** κ**, with and without paranoia scores of zero.** Standardized residual plots (left) and normal probability plots (right). **a,** All 72 subjects in online version 3: a significant regression equation was found (F(3,68)=3.681, p=0.016), with an R^2^ of 0.140. Participants’ predicted κ was equal to 0.486 + 0.062 (PARANOIA)+0.012 (BDI) -0.006 (BAI). Paranoia was a significant predictor of κ (β=0.343, t=2.470, p=0.016, CI=[0.012, 0.113]) but depression and anxiety were not (BDI: β=0.086, t=0.423, p=0.674, CI=[-0.043, 0.066]; BAI: β=-0.043, t=-0.218, p=0.828, CI=[-0.063, 0.050]). **b,** Participants with paranoia scores > 0 (n=39): a significant regression equation was found (F(3,35)=6.322, p=0.002), with an R^2^ of 0.351. Participants’ predicted κ was equal to 0.432 + 0.150 (PARANOIA)+0.013 (BDI) -0.004 (BAI). Paranoia was a significant predictor of κ (β=0.538, t=2.983, p=0.005, CI=[0.048, 0.252]) but depression and anxiety were not (BDI: β=0.111, t=0.494, p=0.624, CI=[-0.041, 0.067]; BAI: β=-0.035, t=-0.163, p=0.872, CI=[-0.057, 0.049]).

**a**

**b**

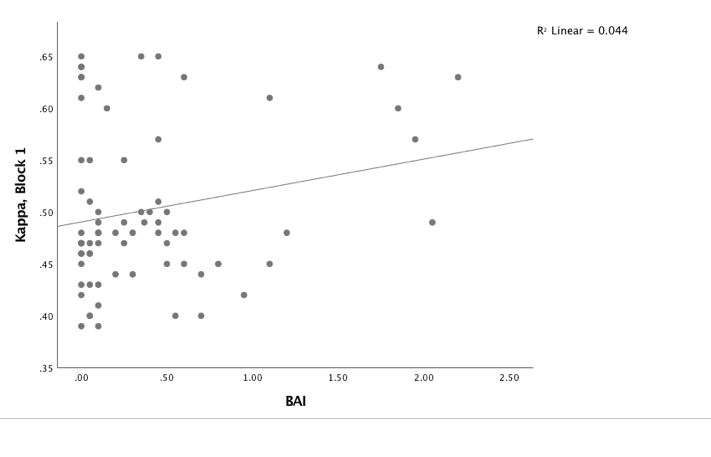

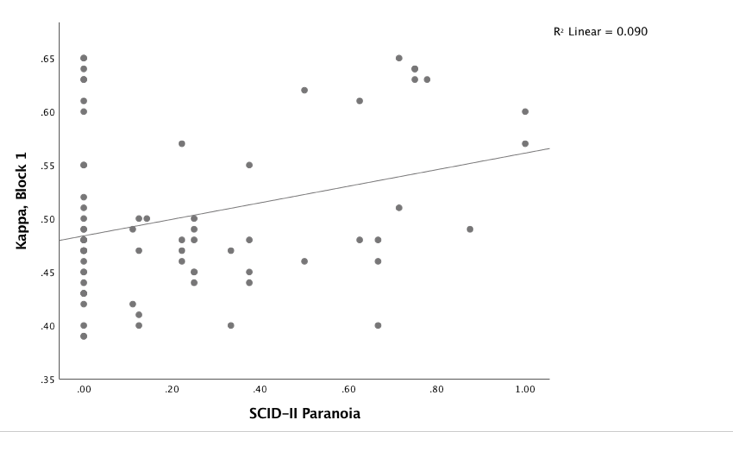

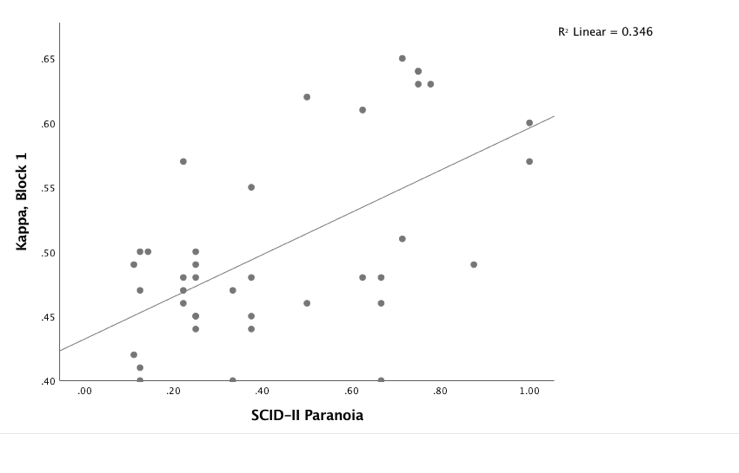

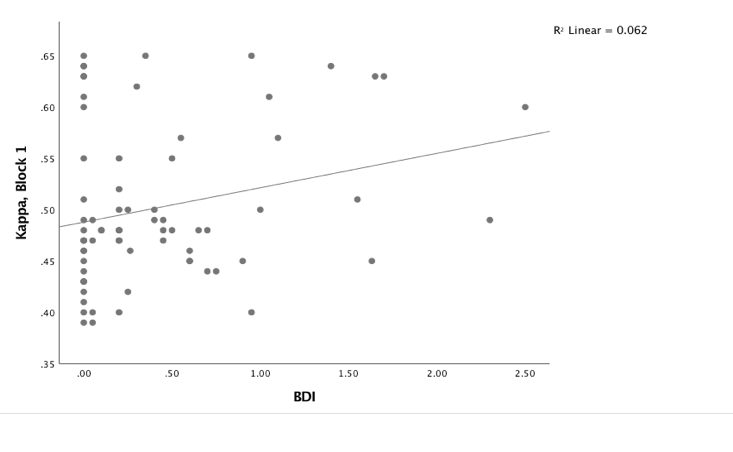

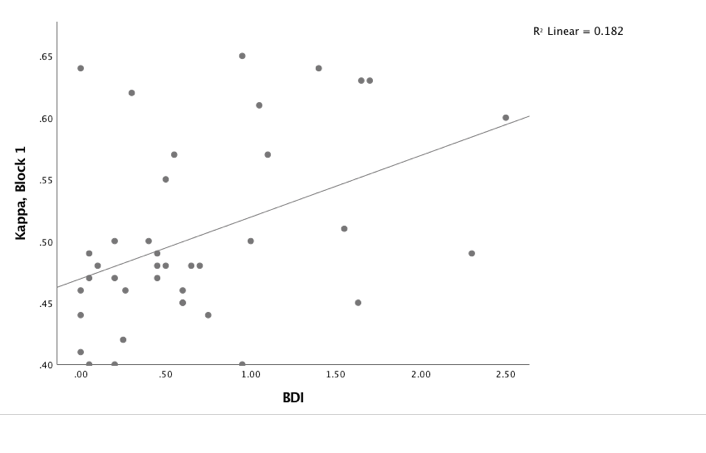

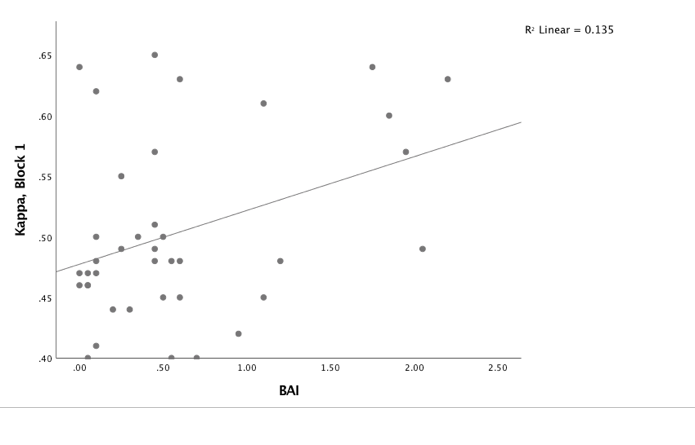

**Supplementary Fig. 2. Correlations between** κ **and symptoms, with and without paranoia scores of zero.** Paranoia (SCID-II, top), depression (BDI, middle), and anxiety (BAI, bottom). **a,** Among all 72 subjects from online version 3, κ correlates with paranoia (r=0.30, p=0.011) and depression (r=0.250, p=0.034), but not anxiety (r=0.210, p=0.077). **b,** Among participants who endorse at least one paranoia item (SCID-II paranoia > 0, n=39), κ correlates with paranoia (r=0.588, p=8.1E-5), depression (r=0.427, p=0.007), and anxiety (r=0.367, p=0.021). All correlations are two-tailed.

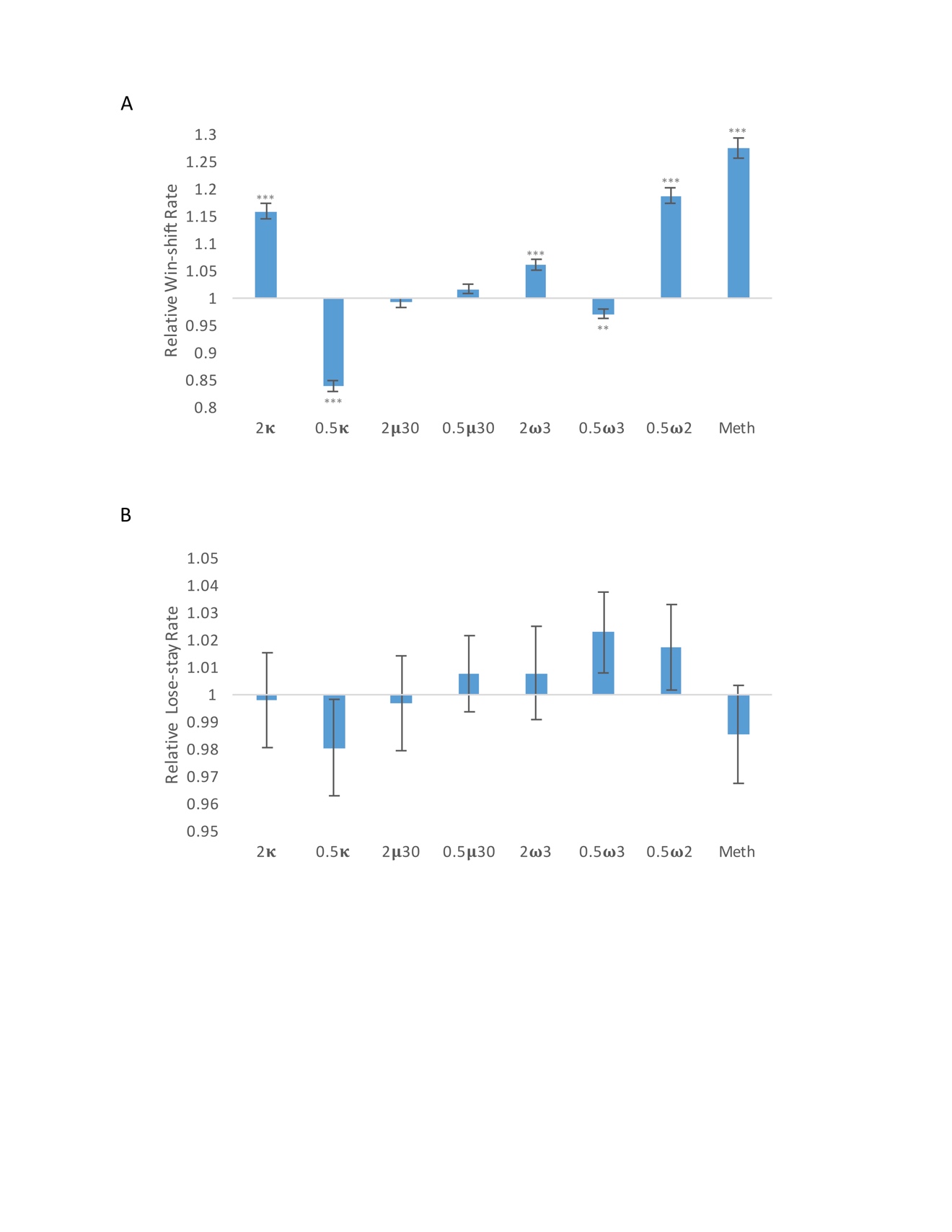

**Supplementary Fig. 3. Parameter effects on simulated task performance.** We simulated behavior from low paranoia participants (online Version 3, n=54) to evaluate the effects of 𝛋, 𝛍_3_^0^, 𝛚_2,_ and 𝛚_3_ on win-shift and lose-stay rates. Estimated perceptual parameters were averaged across subjects to create a single set of baseline parameters. Additional parameter sets were created by doubling or halving one parameter at a time (e.g., 2 𝛋 or 0.5 𝛋), while the others were held constant (n.b., 2 𝛚_2_ violated model assumptions and was excluded from analysis). We also included the average parameter values of rats exposed to methamphetamine (Meth). Ten simulations were run per subject for each condition (i.e., parameter set). Win-shift and lose-stay rates were calculated, then averaged across simulations and subjects. Rates from each condition were divided by the baseline condition rate to generate relative win-shift and lose-stay rates. We compared relative rates for each condition to the baseline (relative rate of 1; paired t-tests, Bonferroni-corrected p-values). Of note, baseline parameters were positive for 𝛋 and 𝛚_2,_ and negative for 𝛍_3_^0^ and 𝛚_3_. Consequently, the doubled (2x) condition makes 𝛍_3_^0^ and 𝛚_3_ more negative (lower). (n=54). Error bars denote standard error (SEM); *p ≤ 0.05, **p ≤ 0.01, ***p ≤ 0.001.

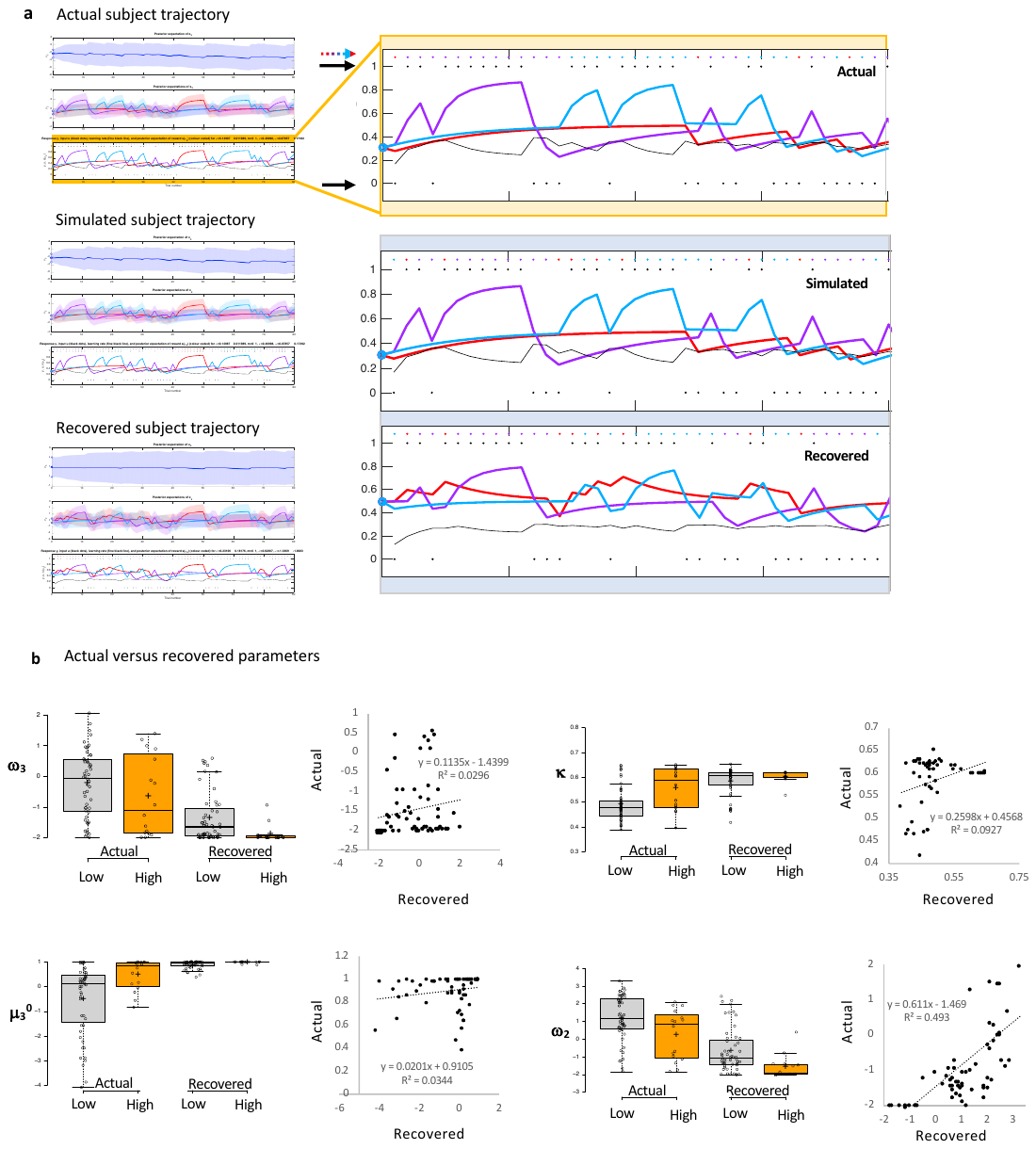

**Supplementary Fig. 4. Parameter recovery.** **a,** Example choice trajectory (top), trajectory simulated from estimated perceptual parameters (middle), and recovered trajectory (bottom). Simulated trajectories closely align with real trajectories, although trial-by-trial choices (colored dots/arrow) occasionally differ. Outcomes (1 or 0; black dots and arrow) remain the same. Recovered trajectories lose some information from the original. **b,** relative to the actual parameters, recovered parameters shift toward configuration file values but maintain some resemblance to original group differences. Original and recovered values significantly correlate for 𝛚_2_ (r=0.702, p=2.52E-11) and 𝛋 (r=0.305, p=0.011) but not 𝛚_3_ (r=0.172, p=0.16) or 𝛍_3_^0^ (r=0.186, p=0.13). Box plots: gray indicates low paranoia, orange designates high paranoia; centre lines depict medians; box limits indicate the 25th and 75th percentiles; whiskers extend 1.5 times the interquartile range from the 25th and 75th percentiles, outliers are represented by dots; crosses represent sample means; data points are plotted as open circles. Online version 3 dataset.

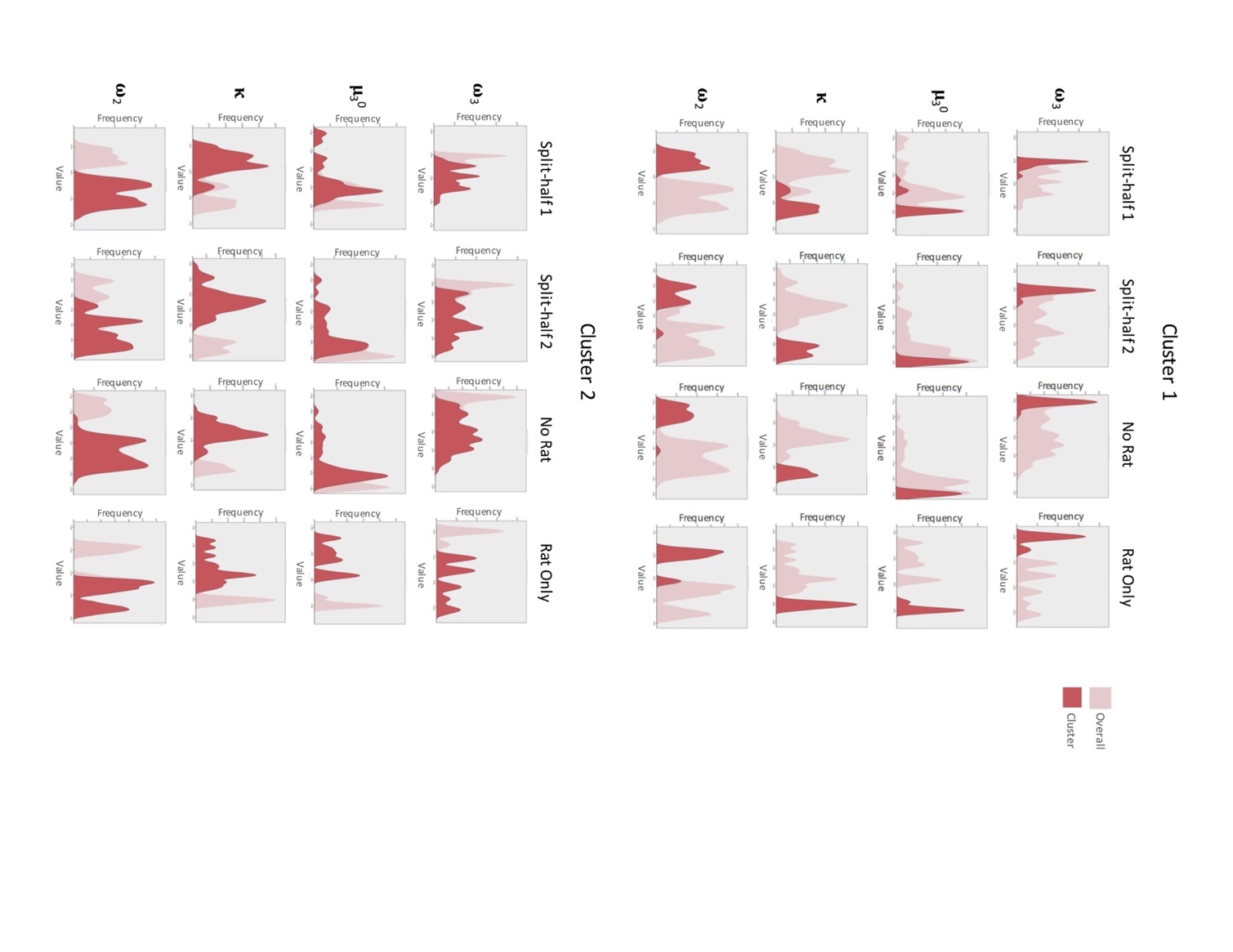

Cluster 2

Cluster 1

**Supplementary Fig. 5. Cluster validation.** We replicated our 2-cluster solution (Fig. 4) by independently running two-step cluster analyses on separate halves of the data (Split-half 1, Split-half 2), removing the rat data and running the human data only (No Rat), and running the rat data alone (Rat Only). In each condition, we identified two clusters with good cohesion and separation (Split-half 1, n=19 cluster 1, 42 cluster 2: silhouette coefficient = 0.6; Split-half 2, n = 17 cluster 1, 43 cluster 2: silhouette coefficient = 0.7; No Rat, n=26 cluster 1, 78 cluster 2: silhouette coefficient = 0.7; Rat Only, n=6 cluster 1, 11 cluster 2: silhouette coefficient = 0.7). All variables showed predictor importance above 0.2 with some variation in order of importance (Split-half 1, 𝛚_2_ > 𝛋 > 𝛚_3_ > 𝛍_3_^0^; Split-half 2, 𝛋 > 𝛚_2_ > 𝛚_3_ > 𝛍_3_^0^; No Rat, 𝛋 > 𝛚_2_ > 𝛚_3_ > 𝛍_3_^0^; Rat Only, 𝛍_3_^0^ > 𝛚_2_ > 𝛚_3_ > 𝛋). Predictor importance was weighted more evenly across variables in the Rat Only condition; all variables showed predictor importance above 0.6.

**a**

| Number of SCID-II Paranoia items endorsed | Number of MTurk Participants |
| --- | --- |
| 0 | 124 |
| 1 | 42 |
| 2 | 39 |
| 3 | 31 |
| 4 | 22 |
| 5 | 15 |
| 6 | 16 |
| 7 | 12 |
| 8 | 5 |
| 9 | 1 |

**bA**

| SCID-II Paranoia Score | Fenigstein & Vanable Score |
| --- | --- |
| 0 | 2.2 |
| 0 | 1.8 |
| 0 | 1.45 |
| 0 | 1.1 |
| 0 | 1.5 |
| 0 | 1.15 |
| 0 | 1.35 |
| 0 | 1.1 |
| 0 | 1.3 |
| 0 | 3.2 |
| 0 | 1.2 |
| 0 | 1.1 |
| 0 | 2.95 |
| 0 | 3.4 |
| 0 | 2.05 |
| 0 | 1.35 |
| 0 | 1.8 |
| 0 | 1.4 |
| 0 | 1.4 |
| 0 | 1.35 |
| 0 | 1.45 |
| 0 | 1 |
| 0.111111 | 1.95 |
| 0.125 | 3.7 |
| 0.125 | 2.25 |
| 0.125 | 2.5 |
| 0.125 | 1.85 |
| 0.125 | 1.6 |
| 0.125 | 4 |
| 0.125 | 1.25 |
| 0.125 | 2 |
| 0.125 | 1.7 |
| 0.222222 | 2.2 |
| 0.25 | 2.35 |
| 0.25 | 2.4 |
| 0.25 | 2.5 |
| 0.25 | 2.4 |
| 0.25 | 2.45 |
| 0.25 | 2.25 |
| 0.285714 | 1.25 |
| 0.375 | 3.31579 |
| 0.375 | 2.31579 |
| 0.375 | 3.15 |
| 0.375 | 1.7 |
| 0.375 | 2.7 |
| 0.428571 | 3.6 |
| 0.444444 | 3 |
| 0.5 | 2.25 |
| 0.5 | 3.95 |
| 0.5 | 2.35 |
| 0.5 | 2.45 |
| 0.5 | 4.2 |
| 0.555556 | 1.95 |
| 0.571429 | 2.57895 |
| 0.571429 | 3.15 |
| 0.625 | 2.5 |
| 0.625 | 4.4 |
| 0.666667 | 2.15 |
| 0.777778 | 2.85 |
| 0.777778 | 3.65 |
| 0.875 | 1.9 |
| 0.888889 | 3.4 |
| 1 | 4.31579 |
| 1 | 4.25 |
| 1 | 3.2 |
| 0 | 1.1 |
| 0 | 1.05 |
| 0 | 1.95 |
| 0 | 1 |
| 0 | 1 |
| 0 | 1.6 |
| 0 | 1.1 |
| 0 | 1.5 |
| 0 | 1.55 |
| 0 | 1.5 |
| 0 | 1 |
| 0 | 1.05 |
| 0 | 1.15 |
| 0 | 1.3 |
| 0 | 1 |
| 0 | 1.6 |
| 0 | 2.05 |
| 0 | 1.85 |
| 0 | 1.5 |
| 0 | 1 |
| 0 | 1.5 |
| 0 | 1.15789 |
| 0 | 1.4 |
| 0 | 1.9 |
| 0 | 1.85 |
| 0 | 1.8 |
| 0 | 1.4 |
| 0 | 1.75 |
| 0 | 1.7 |
| 0 | 1.15 |
| 0 | 1.1 |
| 0 | 1.6 |
| 0 | 1.3 |
| 0.111111 | 2 |
| 0.111111 | 1.35 |
| 0.111111 | 1.65 |
| 0.125 | 1.9 |
| 0.125 | 2.55 |
| 0.125 | 1.8 |
| 0.125 | 2.05 |
| 0.125 | 2.55 |
| 0.125 | 1.65 |
| 0.125 | 2.4 |
| 0.125 | 1.3 |
| 0.125 | 2.45 |
| 0.125 | 1.4 |
| 0.125 | 1.7 |
| 0.125 | 1.45 |
| 0.142857 | 1.15 |
| 0.222222 | 3.55 |
| 0.222222 | 2.85 |
| 0.222222 | 3.45 |
| 0.222222 | 1.75 |
| 0.222222 | 3.05 |
| 0.222222 | 2.6 |
| 0.222222 | 2.55 |
| 0.25 | 2.4 |
| 0.25 | 1.5 |
| 0.25 | 1.45 |
| 0.25 | 1.5 |
| 0.25 | 2 |
| 0.25 | 3.2 |
| 0.333333 | 3.75 |
| 0.375 | 1.05556 |
| 0.375 | 2.95 |
| 0.375 | 2.05 |
| 0.375 | 3.5 |
| 0.375 | 1.75 |
| 0.375 | 3.1 |
| 0.444444 | 1.89474 |
| 0.444444 | 2.4 |
| 0.444444 | 1.95 |
| 0.5 | 2.05 |
| 0.5 | 2.45 |
| 0.5 | 2.65 |
| 0.555556 | 2.57895 |
| 0.555556 | 2.5 |
| 0.625 | 3.2 |
| 0.625 | 3.1 |
| 0.625 | 1.45 |
| 0.666667 | 3.05 |
| 0.666667 | 2.45 |
| 0.666667 | 2.7 |
| 0.666667 | 4.1 |
| 0.777778 | 3.3 |
| 0.777778 | 2.3 |
| 0.875 | 3.05 |
| 0 | 1.1 |
| 0 | 1.85 |
| 0 | 1 |
| 0 | 1.7 |
| 0 | 1.8 |
| 0 | 1.35 |
| 0 | 1.45 |
| 0 | 2.35 |
| 0 | 1.78947 |
| 0 | 1.85 |
| 0 | 1.9 |
| 0 | 1.1 |
| 0 | 1.15 |
| 0 | 1.2 |
| 0 | 1 |
| 0 | 1.15 |
| 0 | 1.4 |
| 0 | 1.25 |
| 0 | 1.75 |
| 0 | 1.3 |
| 0 | 1 |
| 0 | 1 |
| 0 | 2.2 |
| 0 | 1.36842 |
| 0 | 1 |
| 0 | 1 |
| 0 | 1.2 |
| 0 | 1.35 |
| 0 | 1.25 |
| 0 | 1.2 |
| 0 | 1.1 |
| 0 | 1.1 |
| 0 | 1.65 |
| 0.111111 | 2.8 |
| 0.111111 | 1 |
| 0.125 | 1 |
| 0.125 | 1.45 |
| 0.125 | 1.4 |
| 0.125 | 1.95 |
| 0.142857 | 4.3 |
| 0.222222 | 1.65 |
| 0.222222 | 1.85 |
| 0.222222 | 1.75 |
| 0.222222 | 1.84211 |
| 0.25 | 3 |
| 0.25 | 2.75 |
| 0.25 | 1.65 |
| 0.25 | 2.55 |

| SCID-II Paranoia Score | High paranoia |
| --- | --- |
| 0.428571 | 3.6 |
| 0.428571 | 2.42105 |
| 0.444444 | 3 |
| 0.444444 | 1.89474 |
| 0.444444 | 2.4 |
| 0.444444 | 1.95 |
| 0.5 | 2.25 |
| 0.5 | 3.95 |
| 0.5 | 2.35 |
| 0.5 | 2.45 |
| 0.5 | 4.2 |
| 0.5 | 2.05 |
| 0.5 | 2.45 |
| 0.5 | 2.65 |
| 0.5 | 1.55 |
| 0.5 | 1.3 |
| 0.5 | 1.25 |
| 0.5 | 2.45 |
| 0.5 | 3.65 |
| 0.5 | 2.2 |
| 0.5 | 2.75 |
| 0.555556 | 1.95 |
| 0.555556 | 2.57895 |
| 0.555556 | 2.5 |
| 0.555556 | 1.65 |
| 0.555556 | 3.05 |
| 0.571429 | 2.57895 |
| 0.571429 | 3.15 |
| 0.625 | 2.5 |
| 0.625 | 4.4 |
| 0.625 | 3.2 |
| 0.625 | 3.1 |
| 0.625 | 1.45 |
| 0.625 | 2.45 |
| 0.625 | 3.95 |
| 0.625 | 3.55 |
| 0.666667 | 2.15 |
| 0.666667 | 3.05 |
| 0.666667 | 2.45 |
| 0.666667 | 2.7 |
| 0.666667 | 4.1 |
| 0.666667 | 2.5 |
| 0.666667 | 3.15 |
| 0.666667 | 4.35 |
| 0.666667 | 3.4 |
| 0.666667 | 1.85 |
| 0.714286 | 2.9 |
| 0.714286 | 3.84211 |
| 0.75 | 3.7 |
| 0.75 | 3.15 |
| 0.75 | 4 |
| 0.75 | 3.2 |
| 0.75 | 3.9 |
| 0.777778 | 2.85 |
| 0.777778 | 3.65 |
| 0.777778 | 3.3 |
| 0.777778 | 2.3 |
| 0.777778 | 3.75 |
| 0.777778 | 3.94737 |
| 0.8 | 3.85 |
| 0.875 | 1.9 |
| 0.875 | 3.05 |
| 0.875 | 3.95 |
| 0.875 | 4.25 |
| 0.875 | 3.95 |
| 0.888889 | 3.4 |
| 1 | 4.31579 |
| 1 | 4.25 |
| 1 | 3.2 |
| 1 | 2.7 |
| 1 | 3.7 |

**Supplementary Fig. 6. Distribution of paranoia scores. a,** Distribution of total items endorsed on SCID-II measure of paranoid personality in online experiment, all versions (n=307 total). 71 participants endorsed 4 or more items. **b,** Normalized SCID-II paranoia and Fenigstein and Vanable paranoia^3^ scores correlate (r=0.693, p=5.85E-45, 2-tailed). Grey indicates low paranoia (<4/9), and orange indicates high paranoia.

| **Supplementary Table 1. ANOVA Results for HGF Parameters** | | | | | | | | |
| --- | --- | --- | --- | --- | --- | --- | --- | --- |
|  | Block Effect ^†^ | | | Group Effect^‡^ | | | Interaction Effect | |
|  | Statistic^§^ | p-value | Statistic^§^ | | p-value | Statistic^§^ | | p-value |
| **Experiment 1** | | | | | | | | |
| 𝛚_3_ | 11.672 (1) | **0.002** | 1.294 (1) | | 0.264 | 6.948 (1) | | **0.013** |
| 𝛍_3_^0^ | 25.904 (1) | **1.809E-5** | 7.063 (1) | | **0.012** | 5.344 (1) | | **0.028** |
| 𝛋 | 7.768 (1) | **0.009** | 7.599 (1) | | **0.010** | 0.003 (1) | | 0.960 |
| 𝛚_2_ | 2.182 (1) | 0.150 | 4.186 (1) | | **0.050** | 0.058 (1) | | 0.811 |
| 𝛍_2_^0^ | 4.831 (1) | **0.036** | 1.261 (1) | | 0.270 | 0.370 (1) | | 0.547 |
| **BIC** | 0.061 (1) | 0.807 | 8.801 (1) | | **0.006** | 1.7 (1) | | 0.202 |
| **Experiment 2, Version 3** | | | | | | | | |
| 𝛚_3_ | 14.932 (1) | **0.0002** | 1.128 (1) | | 0.292 | 1.406 (1) | | 0.240 |
| 𝛍_3_^0^ | 64.651 (1) | **1.54E-11** | 6.366 (1) | | **0.014** | 0.003 (1) | | 0.959 |
| 𝛋 | 15.53 (1) | **0.0002** | 13.521 (1) | | **0.0005** | 0.011 (1) | | 0.916 |
| 𝛚_2_ | 0.027 (1) | 0.869 | 8.70 (1) | | **0.004** | 0.090 (1) | | 0.765 |
| 𝛍_2_^0^ | 11.432 (1) | **0.001** | 0.030 (1) | | 0.864 | 0.203 (1) | | 0.653 |
| **BIC** | 1.110E-5 (1) | 0.997 | 16.336 (1) | | **0.0001** | 1.678 (1) | | 0.199 |
| **Experiment 3: Rats** | |  |  | |  |  | |  |
| 𝛚_3_ | 30.086 (1) | **6.2785E-5** | 4.579 (1) | | **0.049** | 9.058 (1) | | **0.009** |
| 𝛍_3_^0^ | 31.416 (1) | **5.0188E-5** | 8.454 (1) | | **0.011** | 5.159 (1) | | **0.038** |
| 𝛋 | 9.132 (1) | **0.009** | 13.356 (1) | | **0.002** | 2.644 (1) | | 0.125 |
| 𝛚_2_ | 32.192 (1) | **4.4173E-5** | 22.344 (1) | | **0.0003** | 18.454 (1) | | **0.001** |
| 𝛍_2_^0^ | 5.226 (1) | **0.037** | 0.368 (1) | | 0.553 | 2.087 (1) | | 0.169 |
| **BIC** | 5.052 (1) | **0.040** | 1.890 (1) | | 0.189 | 0.331 (1) | | 0.573 |
| ^†^ Block refers to first versus second half in human studies, Pre-Rx vs Post-Rx in rat studies. | | | | | | | | |
| ^‡^ Group refers to low versus high paranoia in humans, saline versus methamphetamine in rats | | | | | | | | |
| ^§^ F-statistic (degrees of freedom); split-plot ANOVA (i.e., repeated measures with between-subjects factor). | | | | | | | | |

| **Supplementary Table 2. Corrections for Multiple Comparisons** | | | | | | |  |  | |
| --- | --- | --- | --- | --- | --- | --- | --- | --- | --- |
|  | | Group Effect ^†^ | | | | Interaction Effect^‡^ | | | |
|  | | Survives Bonferroni?^§^ | Survives FDR? | Critical Value | Benjamini-Hochberg p-value | Survives Bonferroni?^§^ | Survives FDR? | Critical Value | Benjamini-Hochberg p-value |
| **Experiment 1** | |  |  |  |  |  |  |  |  |
| 𝛚_3_ | | N/A | N/A | 0.05 | 0.264 | No | No | 0.0125 | 0.052 |
| 𝛍_3_^0^ | | **Yes** | **Yes** | **0.025** | **0.024** | No | No | 0.025 | 0.056 |
| 𝛋 | | **Yes** | **Yes** | **0.0125** | **0.04** | N/A | N/A | 0.05 | 0.96 |
| 𝛚_2_ | | No | No | 0.0375 | 0.0667 | N/A | N/A | 0.0375 | 1.081 |
| **Experiment 2, Version 3** | | |  |  |  |  |  |  |  |
| 𝛚_3_ | N/A | | N/A | 0.05 | 0.292 | N/A | N/A | 0.0125 | 0.96 |
| 𝛍_3_^0^ | No | | **Yes** | **3.75E-02** | **0.0187** | N/A | N/A | 0.05 | 0.959 |
| 𝛋 | **Yes** | | **Yes** | **0.0125** | **0.002** | N/A | N/A | 0.0375 | 1.221 |
| 𝛚_2_ | **Yes** | | **Yes** | **0.025** | **0.008** | N/A | N/A | 0.025 | 1.53 |
| **Experiment 3: Rats** | | |  |  |  | |  |  |  |
| 𝛚_3_ | No | | **Yes** | **5.00E-02** | **0.049** | **Yes** | **Yes** | **0.025** | **0.018** |
| 𝛍_3_^0^ | **Yes** | | **Yes** | **3.75E-02** | **0.0147** | No | No | 0.0375 | 0.0507 |
| 𝛋 | **Yes** | | **Yes** | **0.025** | **0.004** | N/A | N/A | 0.05 | 0.125 |
| 𝛚_2_ | **Yes** | | **Yes** | **0.0125** | **0.0012** | **Yes** | **Yes** | **0.0125** | **0.004** |
| N/A denotes to p-values that were not significant before corrections. | | | | | |  |  |  | |
| ^†^ Low versus high paranoia in humans, saline versus methamphetamine in rats. | | | | | |  |  |  | |
| ^‡^ Group by time (i.e., first versus second half in human studies, Pre-Rx vs Post-Rx in rat studies). | | | | | | | |  | |
| ^§^ p-value < 0.0125 | | |  |  |  |  |  |  | |

| **Supplementary Table 3. Experiment 2 Effects Across Block, Paranoia Group, and Task Version** | | | | | | | | | | | | | | | | | | | | | |
| --- | --- | --- | --- | --- | --- | --- | --- | --- | --- | --- | --- | --- | --- | --- | --- | --- | --- | --- | --- | --- | --- |
|  | Block | | Group | | | Version | | | Block*Group*  Version | | Group*Version | | | | Block*Group | | | Block*Version | | | |
|  | F  (df)^†^ | p | F  (df)^†^ | | p | F  (df)^†^ | | p | F  (df)^†^ | p | F  (df)^†^ | | p | | F  (df)^†^ | | p | F  (df)^†^ | | p | |
| 𝛚_3_ | 3.722 (1) | 0.055 | 0.499 (1) | | 0.481 | 2.061 (3) | | 0.105 | 0.415 (3) | 0.742 | 1.005 (3) | | 0.391 | | 0.145 (1) | | 0.704 | 7.0155 (3) | | | **1.42E-4** |
| 𝛍_3_^0^ | 288.1 (1) | **1.01E-45** | 2.604 (1) | | 0.108 | 2.321 (3) | | 0.075 | 0.261 (3) | 0.853 | 2.329 (3) | | 0.075 | | 0.281 (1) | | 0.597 | 0.061 (3) | | | 0.98 |
| 𝛋 | 120.9 (1) | **7.65E-24** | 3.602 (1) | | 0.059 | 5.06  (3) | | **0.002** | 0.08 (3) | 0.971 | 4.178 (3) | | **0.006** | | 1.028 (1) | | 0.312 | 2.559 (3) | | | 0.055 |
| 𝛚_2_ | 35.3 (1) | **7.92E-9** | 4.435 (1) | | **0.036** | 4.155 (3) | | **0.007** | 0.166 (3) | 0.919 | 2.809 (3) | | **0.04** | | 2.387 (1) | | 0.123 | 8.697 (3) | | | **1.5E-5** |
| 𝛍_2_^0^ | 71.3 (1) | **1.33E-15** | 0.242 (1) | | 0.623 | 0.616 (3) | | 0.605 | 1.081 (3) | 0.358 | 0.412 (3) | | 0.744 | | 0.057 (1) | | 0.812 | 1.505 (3) | | | 0.213 |
| **BIC** | 56.6 (1) | **6.23E-13** | 8.073 (1) | | **0.005** | 5.385 (3) | | **0.001** | 0.262 (3) | 0.853 | 4.927 (3) | | **0.002** | | 0.451 (1) | | 0.502 | 11.905 (3) | | | **2.19E-07** |
| ^†^ F-statistic (degrees of freedom); split-plot ANOVA (i.e., repeated measures with two between-subjects factors). | | | | | | | | | | | | | | | | | | | | | |

| **Supplementary Table 4. Experiment 2 ANCOVAs** | | | | | | | | | | | | | | |
| --- | --- | --- | --- | --- | --- | --- | --- | --- | --- | --- | --- | --- | --- | --- |
|  | 𝛚3 | | | 𝛍30 | | | 𝛋 | | | 𝛚2 | | | | |
| Effect | df | F | p-value | df | F | p-value | df | F | p-value | df | F | | p-value | |
| **Demographics (age, gender, ethnicity, and race)** | | | | | | | | | | | | | | |
| Block | 1 | 0.328 | 0.568 | 1 | 10.835 | **0.001** | 1 | 3.425 | 0.066 | 1 | | 2.711 | | 0.101 |
| Block * Age | 1 | 0.659 | 0.418 | 1 | 2.035 | 0.155 | 1 | 2.195 | 0.14 | 1 | | 0.212 | | 0.646 |
| Block * Gender | 1 | 0.363 | 0.547 | 1 | 0.105 | 0.746 | 1 | 4.042 | **0.046** | 1 | | 0.096 | | 0.757 |
| Block * Ethnicity | 1 | 0.016 | 0.901 | 1 | 0.042 | 0.837 | 1 | 0.268 | 0.605 | 1 | | 0.024 | | 0.876 |
| Block * Race | 1 | 3.244 | 0.073 | 1 | 0.279 | 0.598 | 1 | 0.082 | 0.775 | 1 | | 1.386 | | 0.24 |
| Block * Paranoia Group | 1 | 0.001 | 0.969 | 1 | 0.162 | 0.687 | 1 | 0.738 | 0.391 | 1 | | 1.189 | | 0.277 |
| Block * Version | 3 | 7.61 | **7.25E-05** | 3 | 0.561 | 0.641 | 3 | 2.568 | 0.055 | 3 | | 8.613 | | **1.97E-05** |
| Block * Paranoia Group * Version | 3 | 0.451 | 0.717 | 3 | 0.135 | 0.939 | 3 | 0.119 | 0.949 | 3 | | 0.1 | | 0.96 |
| Age | 1 | 3.054 | 0.082 | 1 | 2.974 | 0.086 | 1 | 2.101 | 0.149 | 1 | | 2.339 | | 0.128 |
| Gender | 1 | 0.438 | 0.509 | 1 | 0.02 | 0.886 | 1 | 0.005 | 0.941 | 1 | | 0.014 | | 0.905 |
| Ethnicity | 1 | 0.029 | 0.865 | 1 | 0.059 | 0.808 | 1 | 0.087 | 0.768 | 1 | | 0.221 | | 0.639 |
| Race | 1 | 0.072 | 0.789 | 1 | 2.218 | 0.138 | 1 | 0.373 | 0.542 | 1 | | 0.333 | | 0.564 |
| Paranoia Group | 1 | 4.71E-04 | 0.983 | 1 | 0.741 | 0.39 | 1 | 1.795 | 0.182 | 1 | | 3.302 | | 0.071 |
| Version | 3 | 1.845 | 0.14 | 3 | 1.914 | 0.128 | 3 | 4.975 | **0.002** | 3 | | 3.786 | | **0.011** |
| Paranoia Group * Version | 3 | 0.935 | 0.424 | 3 | 1.911 | 0.129 | 3 | 3.599 | **0.014** | 3 | | 1.919 | | 0.127 |
| **Mental health factors (medication usage, diagnostic category, BAI score, and BDI score)** | | | | | | | | | | | | | | |
| Block | 1 | 3.333 | 0.069 | 1 | 95.753 | **3.12E-19** | 1 | 25.498 | **8.78E-07** | 1 | 8.341 | | **0.004** | |
| Block * BAI | 1 | 0.26 | 0.611 | 1 | 1.532 | 0.217 | 1 | 2.852 | 0.093 | 1 | 0.394 | | 0.531 | |
| Block * BDI | 1 | 0.009 | 0.926 | 1 | 0.208 | 0.649 | 1 | 6.55 | **0.011** | 1 | 0.597 | | 0.441 | |
| Block * Medication Usage | 1 | 0.027 | 0.87 | 1 | 1.288 | 0.258 | 1 | 0.691 | 0.407 | 1 | 0.871 | | 0.352 | |
| Block * Diagnostic Category | 1 | 1.366 | 0.244 | 1 | 1.785 | 0.183 | 1 | 0.063 | 0.803 | 1 | 0.208 | | 0.649 | |
| Block * Paranoia Group | 1 | 0.068 | 0.795 | 1 | 0.298 | 0.586 | 1 | 0.298 | 0.586 | 1 | 0.007 | | 0.935 | |
| Block * Version | 3 | 5.872 | **0.001** | 3 | 0.531 | 0.662 | 3 | 0.906 | 0.439 | 3 | 6.16 | | **0.0005** | |
| Block * Paranoia Group * Version | 3 | 1.024 | 0.383 | 3 | 0.869 | 0.458 | 3 | 0.266 | 0.85 | 3 | 0.095 | | 0.963 | |
| BAI | 1 | 1.108 | 0.294 | 1 | 0.012 | 0.913 | 1 | 0.954 | 0.33 | 1 | 0.921 | | 0.338 | |
| BDI | 1 | 0.037 | 0.848 | 1 | 0.574 | 0.449 | 1 | 1.343 | 0.248 | 1 | 2.372 | | 0.125 | |
| Medication Usage | 1 | 0.327 | 0.568 | 1 | 0.058 | 0.81 | 1 | 0.002 | 0.966 | 1 | 0.467 | | 0.495 | |
| Diagnostic Category | 1 | 4.252 | **0.04** | 1 | 0.004 | 0.949 | 1 | 1.443 | 0.231 | 1 | 1.743 | | 0.188 | |
| Paranoia Group | 1 | 0.057 | 0.811 | 1 | 0.233 | 0.63 | 1 | 1.032 | 0.311 | 1 | 1.695 | | 0.194 | |
| Version | 3 | 3.183 | **0.025** | 3 | 2.73 | **0.045** | 3 | 5.274 | **0.002** | 3 | 4.468 | | **0.004** | |
| Paranoia Group * Version | 3 | 0.311 | 0.818 | 3 | 2.307 | 0.077 | 3 | 4.556 | **0.004** | 3 | 3.397 | | **0.019** | |
| **Global cognitive ability (educational attainment, income, and cognitive reflection)** | | | | | | | | | | | | | | |
| Block | 1 | 1.19E-04 | 0.991 | 1 | 51.264 | **7.60E-12** | 1 | 28.675 | **1.83E-07** | 1 | 18.388 | | **2.51E-05** | |
| Block * Education | 1 | 0.603 | 0.438 | 1 | 0.001 | 0.975 | 1 | 0.033 | 0.856 | 1 | 0.258 | | 0.612 | |
| Block * Income | 1 | 1.211 | 0.272 | 1 | 2.874 | 0.091 | 1 | 3.483 | 0.063 | 1 | 2.421 | | 0.121 | |
| Block * Cognitive Reflection | 1 | 1.83 | 0.177 | 1 | 0.709 | 0.401 | 1 | 1.221 | 0.27 | 1 | 4.667 | | **0.032** | |
| Block * Paranoia Group | 1 | 0.005 | 0.946 | 1 | 0.359 | 0.55 | 1 | 0.263 | 0.608 | 1 | 0.885 | | 0.348 | |
| Block * Version | 3 | 8.861 | **1.27E-05** | 3 | 0.182 | 0.909 | 3 | 2.325 | 0.075 | 3 | 8.815 | | **1.35E-05** | |
| Block * Paranoia Group * Version | 3 | 0.826 | 0.48 | 3 | 0.478 | 0.698 | 3 | 0.15 | 0.929 | 3 | 0.3 | | 0.825 | |
| Education | 1 | 0.111 | 0.739 | 1 | 0.578 | 0.448 | 1 | 1.395 | 0.239 | 1 | 0.608 | | 0.436 | |
| Income | 1 | 2.763 | 0.098 | 1 | 1.382 | 0.241 | 1 | 0.055 | 0.814 | 1 | 1.035 | | 0.31 | |
| Cognitive Reflection | 1 | 0.164 | 0.686 | 1 | 12.807 | **0.0004** | 1 | 0.224 | 0.636 | 1 | 0.807 | | 0.37 | |
| Paranoia Group | 1 | 0.069 | 0.793 | 1 | 0.555 | 0.457 | 1 | 2.477 | 0.117 | 1 | 4.715 | | **0.031** | |
| Version | 3 | 2.104 | 0.1 | 3 | 2.55 | 0.056 | 3 | 5.53 | **0.001** | 3 | 3.799 | | **0.011** | |
| Paranoia Group * Version | 3 | 1.288 | 0.279 | 3 | 2.568 | 0.055 | 3 | 4.469 | **0.004** | 3 | 2.793 | | **0.041** | |

| **Supplementary Table 5. Modified Cognitive Reflection Questionnaire Items** | |
| --- | --- |
| Item | Prompt |
| 1 | A folder and a paper clip cost $1.10 in total. The folder costs $1.00 more than the paper clip.  How much does the paper clip cost? |
| 2 | If it takes 5 clerks 5 minutes to review 5 applications, how long would it take 100 clerks to review 100 applications? |
| 3 | In a garden, there is a cluster of weeds. Every day, the cluster doubles in size. If it takes 48 days for the cluster to cover the entire garden, how long would it take for the cluster to cover half of the garden? |

| **Supplementary Table 6. Simulations and behavior** | | | | | | | | | | | | | | | |
| --- | --- | --- | --- | --- | --- | --- | --- | --- | --- | --- | --- | --- | --- | --- | --- |
|  | | Win-switch Rate | | | U-value | | | | | | Lose-stay Rate | | | | |
| Effect | | df | F | p-value | df | | F | | p-value | | df | | F | | p-value |
| **Experiment 1** | | | | | | | | | | | | | | | |
| Block | | 1 | 1.465 | 0.236 | 1 | | 16.999 | | **0.0003** | | 1 | | 1.334 | | 0.257 |
| Block*Paranoia Group | | 1 | 0.602 | 0.444 | 1 | | 2.393 | | 0.132 | | 1 | | 2.575 | | 0.119 |
| Paranoia Group | | 1 | 3.579 | 0.068 | 1 | | 3.312 | | 0.079 | | 1 | | 2.283 | | 0.141 |
| **Experiment 2, Version 3** | | | | | | | | | | | | | | | |
| Block | | 1 | 0.935 | 0.337 | 1 | | 10.153 | | **0.002** | | 1 | | 0.122 | | 0.728 |
| Block*Paranoia Group | | 1 | 0.001 | 0.982 | 1 | | 0.003 | | 0.958 | | 1 | | 1.93 | | 0.169 |
| Paranoia Group | | 1 | 12.698 | **0.001** | 1 | | 19.209 | | **4.03E-05** | | 1 | | 1.095 | | 0.299 |
| **Simulations^†^** | | | | | | | | | | | | | | | |
| Block | | 1 | 0.176 | 0.676 | 1 | | 3.335 | | 0.072 | | 1 | | 5.073 | | **0.027** |
| Block*Paranoia Group | | 1 | 2.039 | 0.158 | 1 | | 2.624 | | 0.11 | | 1 | | 0.036 | | 0.85 |
| Paranoia Group | | 1 | 15.394 | **0.0002** | 1 | | 13.362 | | **0.0005** | | 1 | | 0.042 | | 0.839 |
|  | ^†^Simulated data from experiment 2, Version 3 | | | | |  | |  | |  | |  | |  | |

| **Supplementary Table 7. Questionnaire item completion (% responses)** | | |
| --- | --- | --- |
| Questionnaire / subscale | Experiment 1 | Experiment 2 |
| **Age** | 90.6% | 99.7% |
| **Gender** | 100.0% | 100.0% |
| **Ethnicity** | 100.0% | 100.0% |
| **Race** | 100.0% | 100.0% |
| **Education** | 100.0% | 99.7% |
| **Meds** | 100.0% | 90.6% |
| **Dx** | 100.0% | 94.1% |
| **Income** | N/A | 98.0% |
| **SCID-II Paranoia - all items** | 96.9% | 94.1% |
| SCID-II Paranoia - 1 item missing | 3.1% | 5.5% |
| SCID-II Paranoia - 3 items missing | 0.0% | 0.3% |
| **Cognitive reflection - all items** | N/A | 97.7% |
| **Beck's Anxiety Inventory (BAI) - all items** | 90.6% | 96.7% |
| BAI - 1 item missing | 3.1% | 2.9% |
| BAI - 2 items missing | 6.3% | 0.3% |
| **Beck's Depression Inventory (BDI) - all items** | 100.0% | 99.0% |
| BDI - 1 item missing | 0.0% | 1.0% |
