## Supplementary material for "Expecting the unexpected: the paranoid style of belief updating across species": Table 1

| **Table 1. In Lab vs. Online Version 3** | | | | | | |  |  | |  | |  | |  |
| --- | --- | --- | --- | --- | --- | --- | --- | --- | --- | --- | --- | --- | --- | --- |
|  | In Lab | | | | | |  | Online Version 3 | | | | | | |
|  | Low Paranoia (n=21) | High Paranoia (n=11) | | Statistic | | p-value |  | Low Paranoia (n=56) | | High Paranoia (n=16) | | Statistic | | p-value |
| **Demographics** |  | | | | | |  |  | | | | | | |
| Age (years) | 36.0 [3.2] | 38.9 [3.9] | | -0.531 (27)† | | 0.6 |  | 38.6 [1.6] | | 32.9 [1.7] | | 2.441 (41.842)† | | **0.019^¶^** |
| Gender |  | | | 0.006 (1)‡ | | 1^§^ |  |  | | | | .780 (1)‡ | | 0.410 |
| *% Female* | 71.4% | 72.7% | | n/a | | n/a |  | 50.0% | | 62.5% | | n/a | | n/a |
| *% Male* | 28.6% | 27.3% | | n/a | | n/a |  | 50.0% | | 37.5% | | n/a | | n/a |
| *% Other or not specified* | 0% | 0% | | n/a | | n/a |  | 0% | | 0% | | n/a | | n/a |
| Education |  | | | 4.972 (6)‡ | | 0.638^§^ |  |  | | | | 5.351 (6)‡ | | 0.549^§^ |
| *% High school degree or equivalent* | 19.0% | 45.5% | | n/a | | n/a |  | 16.1% | | 6.3% | | n/a | | n/a |
| *% Some college or university, no degree* | 14.3% | 0% | | n/a | | n/a |  | 17.9% | | 25.0% | | n/a | | n/a |
| *% Associate degree* | 9.5% | 9.1% | | n/a | | n/a |  | 12.5% | | 12.5% | | n/a | | n/a |
| *% Bachelor's degree* | 23.8% | 27.3% | | n/a | | n/a |  | 35.7% | | 56.3% | | n/a | | n/a |
| *% Master's degree* | 9.5% | 0% | | n/a | | n/a |  | 14.3% | | 0% | | n/a | | n/a |
| *% Doctorate or professional degree* | 4.8% | 0% | | n/a | | n/a |  | 1.8% | | 0% | | n/a | | n/a |
| *% Completed some postgraduate* | 0% | 0% | | n/a | | n/a |  | 1.8% | | 0% | | n/a | | n/a |
| *% Other / not specified* | 19.0% | 18.2% | | n/a | | n/a |  | 0% | | 0% | | n/a | | n/a |
| Ethnicity |  | | | .134 (1)‡ | | 1^§^ |  |  | | | | .117 (1)‡ | | 1^§^ |
| *% Hispanic, Latino, or Spanish origin* | 23.8% | 18.2% | | n/a | | n/a |  | 8.9% | | 6.3% | | n/a | | n/a |
| *% Not of Hispanic, Latino, or Spanish origin* | 76.2% | 81.8% | | n/a | | n/a |  | 91.1% | | 93.8% | | n/a | | n/a |
| Race |  | | | 6.250 (4)‡ | | 0.186^§^ |  |  | | | | 5.368 (4)‡ | | 0.229^§^ |
| *% White* | *61.9%* | *36.4%* | | n/a | | n/a |  | 85.7% | | 75.0% | | n/a | | n/a |
| *% Black or African American* | *19.0%* | *36.4%* | | n/a | | n/a |  | 0% | | 12.5% | | n/a | | n/a |
| *% Asian* | *14.3%* | *9.1%* | | n/a | | n/a |  | 3.6% | | 6.3% | | n/a | | n/a |
| *% American Indian or Alaska Native* | *4.8%* | *0%* | | n/a | | n/a |  | 1.8% | | 6.3% | | n/a | | n/a |
| *% Multiracial* | *0%* | *0%* | | n/a | | n/a |  | 3.6% | | 0% | | n/a | | n/a |
| *% Other / not specified* | *0%* | *18.2%* | | n/a | | n/a |  | 5.4% | | 0% | | n/a | | n/a |
| **Mental Health** |  | | | | | |  |  | | | | | | |
| Psychiatric diagnosis |  | | | 12.329 (2)‡ | | **0.002^§^** |  |  | | | | 7.850 (3)‡ | | **0.039^§^** |
| *% No psychiatric diagnosis* | 71.4% | 9.1% | | adj. residuals | | **0.004** |  | 71.4% | | 50.0% | | adj. residuals | | 0.465 |
| *% Schizophrenia spectrum* | 19.0% | 36.4% | | adj. residuals | | 0.546 |  | 0% | | 6.3% | | adj. residuals | | 0.307 |
| *% Mood disorder* | 9.5% | 54.5% | | adj. residuals | | 0.020^#^ |  | 21.4% | | 43.8% | | adj. residuals | | 0.356 |
| *% Not specified* | 0% | 0% | | adj. residuals | | n/a |  | 7.1% | | 0% | | adj. residuals | | 0.751 |
| % Medicated | 23.8% | 81.8% | | 9.871 (1)‡ | | **0.003^§^** |  | 7.1% | | 31.3% | | 8.730 (2)‡ | | **0.023^§^** |
| Beck's Anxiety Inventory | 0.27 [0.08] | 0.85 [0.17] | | -3.453 (30)† | | **0.002** |  | 0.24 [0.04] | | 0.90 [0.20] | | -3.303 (16.179)† | | **0.004^¶^** |
| Beck's Depression Inventory | 0.23 [0.05] | 0.66 [0.15] | | -2.67 (11.854)† | | **0.021**^¶^ |  | 0.25 [0.04] | | 1.03 [0.19] | | -3.951 (16.659)† | | **0.001^¶^** |
| SCID Paranoia Personality Score | 0.09 [0.02] | 0.63 [0.04] | | -13.476 (30)† | | **2.92E-14** |  | 0.1 [0.02] | | 0.72 [0.04] | | -16.551 (70)† | | **6.712E-26** |
| **Reversal Learning Performance** |  | | | | | |  |  | |  | |  | |  |
| Total points earned | 7061.9 [286.9] | 6290.9 [372.2] | | 1.608 (30)† | | 0.118 |  | 7533.0 [143.8] | | 6503.1 [340.6] | | 3.177 (70)† | | **0.002** |
| Total reversals achieved | 4.8 [0.7] | 2.5 [0.8] | | 2.145 (30)† | | **0.04** |  | 6.3 [0.3] | | 4.9 [0.8] | | 1.758 (20.14)† | | 0.094^¶^ |
| % Achieving reversals | 90.5% | 72.7% | | 1.407 (1)‡ | | 0.327^§^ |  | 100% | | 87.5% | | 7.200 (1)‡ | | **0.047^§^** |
| *Trials to first reversal* | 29.2 [4.5] | 27.9 [11] | | 0.136 (25)† | | 0.893 |  | 20.0 [1.7] | | 13.7 [1.8] | | 1.774 (68)† | | 0.081 |
| % Recovering post-reversal | 81.0% | 54.5% | | 2.490 (1)‡ | | 0.213^§^ |  | 91.1% | | 69.0% | | 3.482 (1)‡ | | 0.097^§^ |
| *Trials to switch* | 1.68 [0.22] | 1.43 [0.20] | | 0.671 (24)† | | 0.509 |  | 2.1 [0.2] | | 2.6 [0.6] | | -1.088 (64)† | | 0.280 |
| *Trials to recovery* | 3.75 [0.51] | 4 [0.93] | | -0.285 (21)† | | 0.779 |  | 2.9 [0.3] | | 4.9 [0.8] | | -2.694 (60)† | | **0.009** |
| Win-switch rate, block 1 (90-50-10) | 0.08 [0.03] | 0.24 [0.09] | | -1.742 (12.379)† | | 0.106^¶^ |  | 0.04 [0.01] | | 0.13 [0.05] | | -1.906 (15.762)† | | 0.075^¶^ |
| Win-switch rate, block 2 (80-40-20) | 0.07 [0.04] | 0.21 [0.1] | | -1.601 (30)† | | 0.12 |  | 0.02 [0.01] | | 0.12 [0.05] | | -2.02 (15.915)† | | 0.061^¶^ |
| Lose-stay rate, block 1 (90-50-10) | 0.19 [0.03] | 0.13 [0.06] | | 0.919 (30)† | | 0.365 |  | 0.30 [0.03] | | 0.39 [0.06] | | -1.425 (70)† | | 0.158 |
| Lose-stay rate, block 2 (80-40-20) | 0.26 [0.05] | 0.12 [0.05] | | 1.817 (30)† | | 0.079 |  | 0.33 [0.03] | | 0.37 [0.06] | | -0.554 (70)† | | 0.581 |
| Null trials | 8.5 [2.8] | 10.4 [3.7] | | -0.391 (30)† | | 0.699 |  | n/a | | n/a | | n/a | | n/a |
| ^†^ Independent samples t-test: t-value (df). Two-tailed p-values reported |  | |  | |  | |  |  |  | |  | |  | |
| ^‡^ Exact test, chi-square coefficient (df) |  | |  | |  | |  |  |  | |  | |  | |
| ^§^ Exact significance (2-sided) |  | |  | |  | |  |  |  | |  | |  | |
| ^¶^ Equal variances not assumed |  | |  | |  | |  |  |  | |  | |  | |
| ^#^ Not significant (bonferonni correction) |  | |  | |  | |  |  |  | |  | |  | |
