## Supplementary material for "Expecting the unexpected: the paranoid style of belief updating across species": Table 2

| **Table 2. Online Experiment** | | | | | | | | | | | | | | | | | | | | | | | | | | | |
| --- | --- | --- | --- | --- | --- | --- | --- | --- | --- | --- | --- | --- | --- | --- | --- | --- | --- | --- | --- | --- | --- | --- | --- | --- | --- | --- | --- |
|  | | | Version 1 | | | | Version 2 | | | | Version 3 | | | | Version 4 | | | | Version Effect | | | | Paranoia Effect | | | Interaction | |
|  | | | Low Paranoia (n=45) | | High Paranoia (n=20) | | Low Paranoia (n=69) | | High Paranoia (n=18) | | Low Paranoia (n=56 ) | | High Paranoia (n=16) | | Low Paranoia (n=64) | | High Paranoia (n=19) | | Statistic | | p-value | | Statistic | | p-value | Statistic | p-value |
| **Demographics** | | |  | | | | | | | | | | | | | | | | | | | | | | | | |
| Age (years) | | | 36.5 [1.5] | | 35.4 [2.4] | | 36.2 [1.4] | | 39.5 [2.8] | | 38.6 [1.6] | | 32.9 [1.7] | | 37.6 [1.3] | | 30.7 [1.6] | | 1.119 (3)^††^ | | 0.342 | | 3.202 (1)^††^ | | 0.075 | 2.619 (3)^††^ | 0.051 |
| Gender | | |  | | | | | | | | | | | | | | | | 7.289 (6)‡ | | 0.238^§^ | | 1.373 (2)‡ | | 0.503^§^ | n/a | n/a |
| *% Female* | | | 44.4% | | 45.0% | | 47.8% | | 50.0% | | 50.0% | | 62.5% | | 57.8% | | 73.7% | | n/a | | n/a | | n/a | | n/a | n/a | n/a |
| *% Male* | | | 55.6% | | 55.0% | | 50.7% | | 50.0% | | 50.0% | | 37.5% | | 42.2% | | 26.3% | | n/a | | n/a | | n/a | | n/a | n/a | n/a |
| *% Other or not specified* | | | 0% | | 0% | | 1.4% | | 0% | | 0% | | 0% | | 0% | | 0% | | n/a | | n/a | | n/a | | n/a | n/a | n/a |
| Education | | |  | | | | | | | | | | | | | | | | 15.943 (21)‡ | | 0.812^\|\|^ | | 7.326 (7)‡ | | 0.4^§^ | n/a | n/a |
| *% High school degree or equivalent* | | | 17.8% | | 20.0% | | 13.0% | | 16.7% | | 16.1% | | 6.3% | | 25.0% | | 10.5% | | n/a | | n/a | | n/a | | n/a | n/a | n/a |
| *% Some college or university, no degree* | | | 22.2% | | 30.0% | | 24.6% | | 22.2% | | 17.9% | | 25.0% | | 25.0% | | 26.3% | | n/a | | n/a | | n/a | | n/a | n/a | n/a |
| *% Associate degree* | | | 13.3% | | 15.0% | | 17.4% | | 22.2% | | 12.5% | | 12.5% | | 9.4% | | 21.1% | | n/a | | n/a | | n/a | | n/a | n/a | n/a |
| *% Bachelor's degree* | | | 33.3% | | 35.0% | | 40.6% | | 22.2% | | 35.7% | | 56.3% | | 28.1% | | 31.6% | | n/a | | n/a | | n/a | | n/a | n/a | n/a |
| *% Master's degree* | | | 8.9% | | 0% | | 2.9% | | 0% | | 14.3% | | 0% | | 7.8% | | 10.5% | | n/a | | n/a | | n/a | | n/a | n/a | n/a |
| *% Doctorate or professional degree* | | | 4.4% | | 0% | | 0% | | 5.6% | | 1.8% | | 0% | | 1.6% | | 0% | | n/a | | n/a | | n/a | | n/a | n/a | n/a |
| *% Completed some postgraduate* | | | 0% | | 0% | | 1.4% | | 5.6% | | 1.8% | | 0% | | 3.1% | | 0% | | n/a | | n/a | | n/a | | n/a | n/a | n/a |
| *% Other / not specified* | | | 0% | | 0% | | 0% | | 5.6% | | 0% | | 0% | | 0% | | 0% | | n/a | | n/a | | n/a | | n/a | n/a | n/a |
| Income | | |  | | | | | | | | | | | | | | | | 14.961 (18)‡ | | .671^\|\|^ | | 1.177 (6)‡ | | 0.981^§^ | n/a | n/a |
| *Less than $20,000* | | | 24.4% | | 25.0% | | 24.6% | | 33.3% | | 17.9% | | 37.5% | | 23.4% | | 15.8% | | n/a | | n/a | | n/a | | n/a | n/a | n/a |
| *$20,000 to $34,999* | | | 40.0% | | 25.0% | | 20.3% | | 22.2% | | 33.9% | | 31.3% | | 28.1% | | 31.6% | | n/a | | n/a | | n/a | | n/a | n/a | n/a |
| *$35,000 to $49,999* | | | 15.6% | | 15.0% | | 18.8% | | 16.7% | | 12.5% | | 6.3% | | 18.8% | | 15.8% | | n/a | | n/a | | n/a | | n/a | n/a | n/a |
| *$50,000 to $74,999* | | | 13.3% | | 35.0% | | 20.3% | | 5.6% | | 21.4% | | 12.5% | | 18.8% | | 21.1% | | n/a | | n/a | | n/a | | n/a | n/a | n/a |
| *$75,000 to $99,999* | | | 4.4% | | 0% | | 7.2% | | 11.1% | | 8.9% | | 6.3% | | 7.8% | | 15.8% | | n/a | | n/a | | n/a | | n/a | n/a | n/a |
| *Over $100,000* | | | 0% | | 0% | | 5.8% | | 5.6% | | 3.6% | | 6.3% | | 1.6% | | 0% | | n/a | | n/a | | n/a | | n/a | n/a | n/a |
| *Not specified* | | | 2.2% | | 0% | | 2.9% | | 5.6% | | 1.8% | | 0% | | 1.6% | | 0% | | n/a | | n/a | | n/a | | n/a | n/a | n/a |
| Cognitive Reflection | | |  | | | | | | | | | | | | | | | | 11.922 (9)‡ | | 0.223^\|\|^ | | 7.002 (3)‡ | | 0.071^§^ | n/a | n/a |
| *% Answering 0/3 correctly* | | | 11.1% | | 25.0% | | 10.1% | | 11.1% | | 17.9% | | 25.0% | | 15.6% | | 26.3% | | n/a | | n/a | | n/a | | n/a | n/a | n/a |
| *% Answering 1/3 correctly* | | | 4.4% | | 5.0% | | 15.9% | | 11.1% | | 8.9% | | 25.0% | | 14.1% | | 15.8% | | n/a | | n/a | | n/a | | n/a | n/a | n/a |
| *% Answering 2/3 correctly* | | | 13.3% | | 25.0% | | 15.9% | | 16.7% | | 19.6% | | 25.0% | | 21.9% | | 31.6% | | n/a | | n/a | | n/a | | n/a | n/a | n/a |
| *% Answering 3/3 correctly* | | | 71.1% | | 45.0% | | 58.0% | | 61.1% | | 53.6% | | 25.0% | | 48.4% | | 26.3% | | n/a | | n/a | | n/a | | n/a | n/a | n/a |
| Ethnicity | | |  | | | | | | | | | | | | | | | | 5.162 (3)‡ | | 0.157^§^ | | 3.715 (1)‡ | | 0.069^§^ | n/a | n/a |
| *% Hispanic, Latino, or Spanish origin* | | | 4.4% | | 15.0% | | 1.4% | | 0% | | 8.9% | | 6.3% | | 1.6% | | 15.8% | | n/a | | n/a | | n/a | | n/a | n/a | n/a |
| *% Not of Hispanic, Latino, or Spanish origin* | | | 95.6% | | 85.0% | | 98.6% | | 100.0% | | 91.1% | | 93.8% | | 98.4% | | 84.2% | | n/a | | n/a | | n/a | | n/a | n/a | n/a |
| Race | | |  | | | | | | | | | | | | | | | | 19.559 (15)‡ | | .173^\|\|^ | | 9.626 (5)‡ | | 0.084^§^ | n/a | n/a |
| *% White* | | | 82.2% | | 75.0% | | 84.1% | | 88.9% | | 85.7% | | 75.0% | | 85.9% | | 73.7% | | n/a | | n/a | | n/a | | n/a | n/a | n/a |
| *% Black or African American* | | | 6.7% | | 15.0% | | 5.8% | | 11.1% | | 0% | | 12.5% | | 4.7% | | 10.5% | | n/a | | n/a | | n/a | | n/a | n/a | n/a |
| *% Asian* | | | 8.9% | | 10.0% | | 7.2% | | 0% | | 3.6% | | 6.3% | | 7.8% | | 0% | | n/a | | n/a | | n/a | | n/a | n/a | n/a |
| *% American Indian or Alaska Native* | | | 0% | | 0% | | 0% | | 0% | | 1.8% | | 6.3% | | 0% | | 0% | | n/a | | n/a | | n/a | | n/a | n/a | n/a |
| *% Multiracial* | | | 2.2% | | 0% | | 1.4% | | 0% | | 3.6% | | 0% | | 1.6% | | 15.8% | | n/a | | n/a | | n/a | | n/a | n/a | n/a |
| *% Other / not specified* | | | 0% | | 0% | | 1.4% | | 0% | | 5.4% | | 0% | | 0% | | 0% | | n/a | | n/a | | n/a | | n/a | n/a | n/a |
| **Mental Health** | | |  | | | | | | | | | | | | | | | | | | | | | | | | |
| Psychiatric diagnosis | | |  | | | | | | | | | | | | | | | | 10.783 (9)‡ | | 0.292^\|\|^ | | 2.960 (3)‡ | | 0.361^§^ | n/a | n/a |
| *% No psychiatric diagnosis* | | | 73.3% | | 80.0% | | 60.9% | | 55.6% | | 71.4% | | 50.0% | | 65.6% | | 42.1% | | n/a | | n/a | | n/a | | n/a | n/a | n/a |
| *% Schizophrenia spectrum* | | | 2.2% | | 0% | | 0% | | 0% | | 0% | | 6.3% | | 0% | | 0% | | n/a | | n/a | | n/a | | n/a | n/a | n/a |
| *% Mood disorder* | | | 13.3% | | 15.0% | | 27.5% | | 22.2% | | 21.4% | | 43.8% | | 26.6% | | 31.6% | | n/a | | n/a | | n/a | | n/a | n/a | n/a |
| *% Not specified* | | | 11.1% | | 5.0% | | 11.6% | | 22.2% | | 7.1% | | 0% | | 7.8% | | 26.3% | | n/a | | n/a | | n/a | | n/a | n/a | n/a |
| % Medicated | | | 8.9% | | 10.0% | | 13.0% | | 22.2% | | 7.1% | | 31.3% | | 14.1% | | 10.5% | | 3.575 (6)‡ | | 0.744^§^ | | 4.164 (2)‡ | | 0.121^§^ | n/a | n/a |
| Beck's Anxiety Inventory | | | 0.34 [0.06] | | 0.52 [0.14] | | 0.31 [0.04] | | 0.6 [0.13] | | 0.24 [0.04] | | 0.90 [0.20] | | 0.33 [0.06] | | 0.79 [0.18] | | 1.244 (3)^††^ | | 0.2941 | | 38.752 (1)^††^ | | **1.63E-09** | 2.577 (3)^††^ | 0.0539 |
| Beck's Depression Inventory | | | 0.36 [0.07] | | 0.86 [0.15] | | 0.32 [0.05] | | 0.79 [0.13] | | 0.25 [0.04] | | 1.03 [0.19] | | 0.38 [0.07] | | 1.06 [0.20] | | 1.023 (3)^††^ | | 0.3827 | | 74.528 (1)^††^ | | **3.62E-16** | 1.089 (3)^††^ | 0.3542 |
| SCID Paranoia Personality Score | | | 0.11 [0.02] | | 0.67 [0.04] | | 0.11 [0.02] | | 0.61 [0.03] | | 0.1 [0.02] | | 0.72 [0.04] | | 0.11 [0.02] | | 0.65 [0.03] | | 1.297 (3)^††^ | | 0.2756 | | 879.379 (1)^††^ | | **4.81E-91** | 2.018 (3)^††^ | 0.1114 |
| **Reversal Learning Performance** | | |  | | | | | | | | | | | | | | | | | | | | | | |  |  |
| Total points earned | | | 8656.7 [182.9] | | 8372.5 [405.2] | | 6045.7 [135.7] | | 6266.7 [288.0] | | 7533.0 [143.8] | | 6503.1 [340.6] | | 7171.1 [175.6] | | 6510.5 [403.6] | | 32.288 (3)^††^ | | **4.16E-18** | | 6.175 (1)^††^ | | **0.0135** | 2.258 (3)^††^ | 0.0818 |
| Total reversals achieved | | | 7.2 [0.3] | | 6.5 [0.5] | | 5.5 [0.3] | | 5.7 [0.5] | | 6.3 [0.3] | | 4.9 [0.8] | | 5.9 [0.3] | | 4.8 [0.6] | | 4.329 (3)^††^ | | **0.005** | | 5.762 (1)^††^ | | **0.017** | 1.101 (3)^††^ | 0.349 |
| % Achieving reversals | | | 100% | | 100% | | 98.6% | | 94.4% | | 100% | | 87.5% | | 96.9% | | 94.7% | | 2.26 (3)‡ | | 0.598^§^ | | 4.4 (1)‡ | | 0.058^§^ | n/a | n/a |
| Win-switch rate, block 1 (90-50-10) | | | 0.09 [0.03] | | 0.09 [0.04] | | 0.07 [0.01] | | 0.11 [0.05] | | 0.04 [0.01] | | 0.13 [0.05] | | 0.1 [0.03] | | 0.21 [0.06] | | 2.284 (3)^††^ | | 0.079 | | 7.117 (1)^††^ | | **0.008** | 1.15 (3)^††^ | 0.329 |
| Win-switch rate, block 2 (80-40-20) | | | 0.05 [0.02] | | 0.08 [0.03] | | 0.04 [0.01] | | 0.05 [0.04] | | 0.02 [0.01] | | 0.12 [0.05] | | 0.06 [0.02] | | 0.15 [0.05] | | 2.067 (3)^††^ | | 0.105 | | 9.918 (1)^††^ | | **0.002** | 1.174 (3)^††^ | 0.32 |
| Lose-stay rate, block 1 (90-50-10) | | | 0.27 [0.03] | | 0.34 [0.05] | | 0.37 [0.03] | | 0.34 [0.04] | | 0.3 [0.03] | | 0.39 [0.06] | | 0.32 [0.03] | | 0.34 [0.04] | | 0.561 (3)^††^ | | 0.641 | | 1.834 (1)^††^ | | 0.177 | 0.754 (3)^††^ | 0.521 |
| Lose-stay rate, block 2 (80-40-20) | | | 0.28 [0.03] | | 0.23 [0.05] | | 0.4 [0.03] | | 0.32 [0.05] | | 0.33 [0.03] | | 0.37 [0.06] | | 0.29 [0.03] | | 0.33 [0.06] | | 2.47 (3)^††^ | | 0.062 | | 0.177 (1)^††^ | | 0.674 | 0.834 (3)^††^ | 0.476 |
| Reaction time, block 1 | | | 433.6 [28.8] | | 789.3 [282.7] | | 548.1 [77.8] | | 365.6 [26.4] | | 448 [60.1] | | 442.1 [59.5] | | 557.2 [108.2] | | 530 [130.2] | | 0.793 (3)^††^ | | 0.499 | | 0.161 (1)^††^ | | 0.689 | 1.727 (3)^††^ | 0.161 |
| Reaction time, block 2 | | | 370.7 [23.3] | | 494.3 [88.6] | | 465.3 [61.6] | | 331.4 [22.9] | | 391.7 [52.3] | | 555.9 [121.2] | | 385.4 [29.2] | | 504.1 [82.7] | | 0.394 (3)^††^ | | 0.757 | | 1.92 (1)^††^ | | 0.167 | 1.949 (3)^††^ | 0.122 |
| ^††^Univariate analysis, F(df)  ^‡^ Exact test, chi-square coefficient (df)  ^§^ Exact significance (2-sided) | | |  | |  | |  | |  | |  | |  | |  | |  | |  | |  | |  | |  |  |  |
| ^\|\|^ Monte Carlo significance (2-sided) | | |  | |  | |  | |  | |  | |  | |  | |  | |  | |  | |  | |  |  |  |
