## Supplementary material for "Expecting the unexpected: the paranoid style of belief updating across species": Table 3

| **Table 3. Summary of Paranoia / Methamphetamine Effects on Belief-Updating** | | | |
| --- | --- | --- | --- |
|  | In lab | Online | Rats |
| 𝛚_3_ | ↓^†^ | ⇣ | ⬇ |
| 𝛍_3_^0^ | ⬆ | ⬆^‡§^ | ⬆ |
| 𝛋 | ⬆ | ⬆^‡^ | ⬆ |
| 𝛚_2_ | ⬇ | ⬇^‡¶^ | ⬇ |
| 𝛍_2_^0^ | - | - | - |
| ⇡⇣ Non-significant increase/decrease in high paranoia or meth, relative to low paranoia or saline | | | |
| ↑ ↓ Trend-level increase/decrease in high paranoia or meth, relative to low paranoia or saline | | | |
| **⬆⬇** Significantly higher/lower in high paranoia or meth, relative to low paranoia or saline | | | |
| - - No significant findings or trends | |  |  |
| ^†^Baseline trend; parameter decreases in second block for low but not high paranoia | | | |
| ^‡^Version 3 only |  |  |  |
| ^§^Trend-level significance disappears with inclusion of demographic covariates | | | |
| ^¶^ Significance reduced to trend with inclusion of demographic covariates | | |  |
